# Noise in neurons and synapses enables reliable associative memory storage in local cortical circuits

**DOI:** 10.1101/583922

**Authors:** Chi Zhang, Danke Zhang, Armen Stepanyants

## Abstract

Neural networks in the brain can function reliably despite various sources of errors and noise present at every step of signal transmission. These sources include errors in the presynaptic inputs to the neurons, noise in synaptic transmission, and fluctuations in the neurons’ postsynaptic potentials. Collectively they lead to errors in the neurons’ outputs which are, in turn, injected into the network. Does unreliable network activity hinder fundamental functions of the brain, such as learning and memory retrieval? To explore this question, this article examines the effects of errors and noise on properties of biologically constrained networks of inhibitory and excitatory neurons involved in associative sequence learning. The associative learning problem is solved analytically and numerically, and it is also shown how memory sequences can be loaded into the network with a more biologically plausible perceptron-type learning rule. Interestingly, the results reveal that errors and noise during learning increase the probability of memory recall. There is a tradeoff between the capacity and reliability of stored memories, and, noise during learning is required for optimal retrieval of stored information. What is more, networks loaded with associative memories to capacity display many structural and dynamical features observed in local cortical circuits. Due to the similarities between the associative and brain networks, this article predicts that the connections originating from unreliable inhibitory and excitatory neurons or neuron classes in the cortex must be depressed or eliminated during learning, while the connections onto noisy neurons or neuron classes must have lower probabilities and higher weights.

## INTRODUCTION

Brain networks can reliably store and retrieve long-term memories despite the facts that various sources of errors and noise accompany every step of signal transmission through the network (1), synaptic connectivity changes over time (2-4), and extraneous sensory inputs are usually present during memory recall. The brain can reduce the effects of noise and extraneous sensory inputs by attending to the memory retrieval process (5, 6), but such hinderances cannot be eliminated entirely. Therefore, the reliability required for memory retrieval must be built into the network during learning. This proposal presents an interesting challenge. Traditional supervised learning models, such as the ones that rely on the perceptron rule (7, 8), modify connectivity only when a neuron’s output deviates from its target output. Thus, in such models learning stops as soon as the neuron produces the desired response and, subsequently, there is no possibility for improving the response reliability. The network connection weights in such models may end up near the boundary of the solution region, and a small amount of noise or extraneous input during memory retrieval can lead to errors or completely disrupt the retrieval process. More reliable solutions are located farther away from the solution region boundary, but the perceptron rule is not guaranteed to find them. Thus, it is not clear how the neural networks in the brain manage not only to learn but also to do it reliably.

In the case of associative memory storage, reliability can be incorporated into the perceptron learning rule by means of a generic robustness parameter [see e.g. (9, 10)]. This simplified description, however, is neither biologically motivated nor does it reflect the recurrent cycle of errors and noise present during learning and memory retrieval (Figure 1A). A more comprehensive account must include errors in the input to the neurons, combine them with fluctuations in the neurons’ presynaptic weights and intrinsic sources of noise, and produce spiking errors in the neurons’ outputs. The latter injected back into the network, would give rise to input errors for the next time step, completing the error propagation cycle. The recurrence of errors presents a clear challenge for the retrieval of associative memory sequences considered in this study. If not corrected at every step of the retrieval process, errors in the network activity can amplify over time and lead to an irreversible deviation of the retrieved trajectory from the loaded sequence, i.e. a partially retrieved memory.

**Figure 1:**
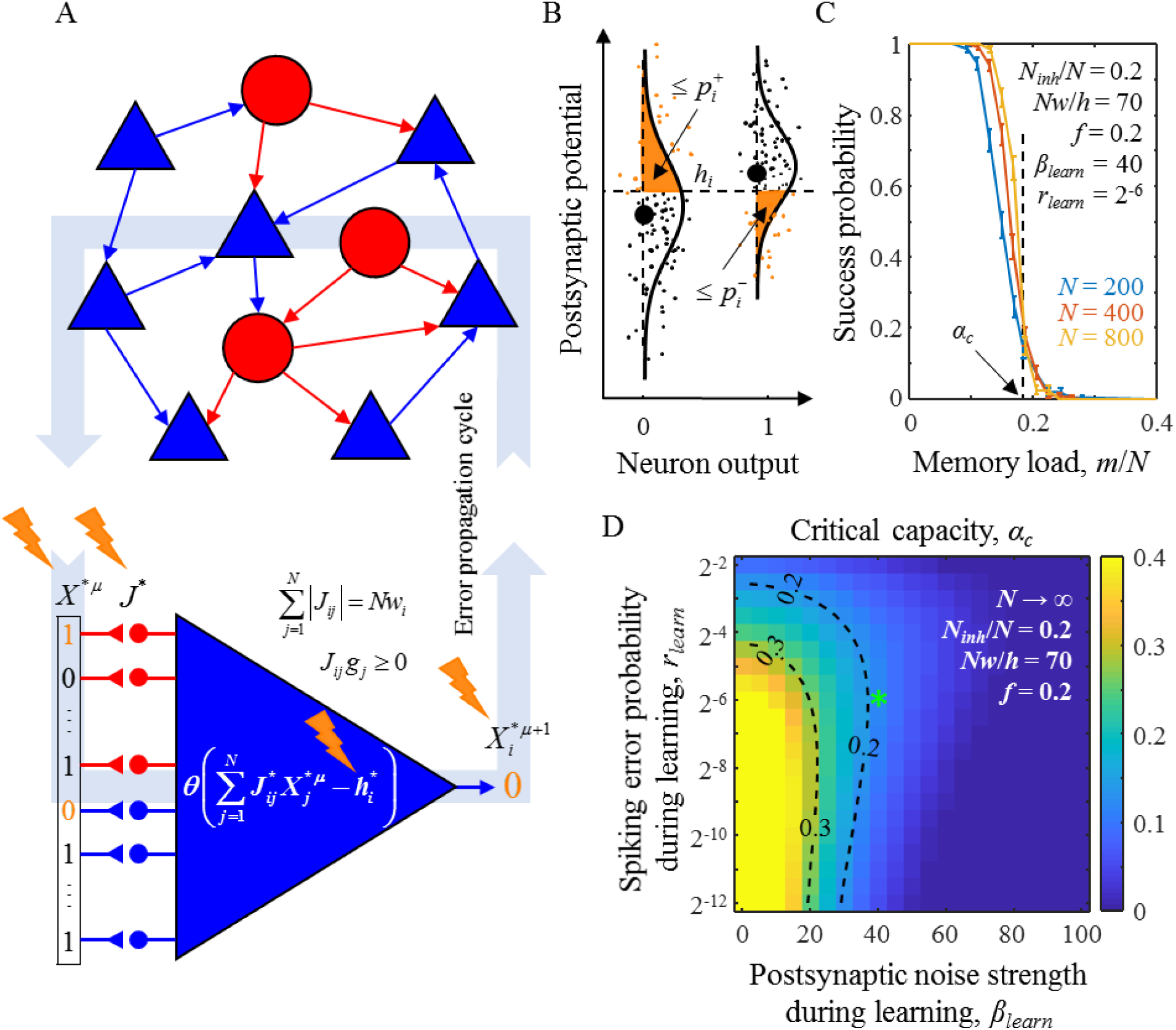
Associative memory storage in a recurrent network of inhibitory and excitatory neurons in the presence of errors and noise. **A.** A cycle of error propagation through the network. Inhibitory neurons (red circles) and excitatory neurons (blue triangles) form an all-to-all potentially (structurally) connected network. Red and blue arrows represent actual (functional) connections. Spiking errors, *X*^*^, synaptic noise, 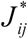, and intrinsic noise, 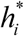, accompany signal transmission (orange lightning signs). Errors in the neurons’ outputs at a given time step become the spiking errors for the next time step. **B.** Fluctuations in postsynaptic potentials for two associations with target neuron outputs 0 (left) and 1 (right). Large black dots denote postsynaptic potentials in the absence of errors and noise. Small dots represent postsynaptic potentials on different trials in the presence of errors and noise. Orange areas to the left of the postsynaptic potential probability densities (solid lines) represent the probabilities of erroneous spikes (left) and spike failures (right). **C.** The probability of successful learning by a neuron is a sharply decreasing function of associative sequence length, *m* (or memory load *m*/*N*). Solid curves represent the probabilities of successful learning obtained with nonlinear optimization (see section I of SI) for neurons receiving *N* = 200, 400, and 800 homogeneous inputs (same *f* and *r*). The values of *β*_*learn*_ and *r*_*learn*_ are provided in the figure. The numerical values of all other parameters of the model were adapted from (27). At 0.5 success probability, the neuron is said to be loaded to capacity, *α*. The dashed black line represents the theoretical (critical) capacity, *α*_*c*_, obtained with the replica method in the *N* → ∞ limit. **D.** Map of *α*_*c*_ for a neuron receiving homogeneous input as a function of *β*_*learn*_ and *r*_*learn*_. The map was obtained with the replica method (see sections A-H of SI). The green asterisk indicates the values of *β*_*learn*_ and *r*_*learn*_ used in (C). Dashed isocontours are drawn as a guide to the eye.

The premise of this article is that errors and noise are essential components of the reliable learning mechanism implemented in the brain. As different fluctuations accompany the presentation of the same learning example to a neuron on different trials, the neuron in effect never stops learning. Its connection weights move further away from the solution region boundary every time a progressively larger fluctuation is encountered. This process increases the reliability of the loaded memory which can later be retrieved in the presence of noise. Similar ideas have been successfully used in machine learning where an augmentation of training examples with noise (11) and dropping out neurons and connections (12) during training have been shown to significantly reduce both overfitting and training time. There are many other examples in which noise is put to constructive use to improve various functions of physical and neural systems [for reviews see (13-16)]. Therefore, the hypothesis that errors and noise are exploited by the brain for reliable memory storage is not entirely surprising. Still, this hypothesis requires careful quantitative evaluation and validation with experimental data which is the focus of this study.

To that end, we examined a network model of biologically constrained inhibitory and excitatory McCulloch and Pitts neurons (17) loaded to capacity with associative memory sequences (18) in the presence of errors and noise. First, we show that there is a tradeoff between the capacity and reliability of stored sequences which is controlled by the levels of errors and noise present during learning. For optimal tradeoff, as judged by the amount of information contained in the retrieved sequences, noise must be present during learning. Second, as synaptic connectivity of neurons changes during learning (3), it is not unreasonable to expect that the requirement of reliable memory retrieval is reflected in the properties of network connectivity and, consequently, the activity of neurons in the brain. Interestingly, local neural networks in the mammalian cortical areas have many common features of connectivity and network activity (18). We show that these network properties in the model emerge all at once during reliable memory storage. Third, as levels of errors and noise can differ across individual neurons or neuron classes, we examined the properties of model networks composed of heterogeneous neurons and make two salient predictions regarding the connectivity of noisy neurons.

We would like to mention other studies that incorporated sources of noise into the associative learning model and examined the effects of learning on neural network properties. In these studies, the basic associative learning model (7, 19-24) was extended to include biologically inspired elements, such as sign-constrained postsynaptic connections (inhibitory and excitatory) [see e.g. (9, 25, 26)], homeostatically constrained presynaptic connections [see e.g. (27)], and robustness to noise [see e.g. (22, 23)]. In particular, Brunel et al. (9, 28) calculated neuron capacity in the presence of intrinsic noise and output spiking errors [see Supplementary Material in (9)] and showed that sparse excitatory connectivity and certain two- and three-neuron motifs develop in networks loaded to capacity. Rubin et al. (10) considered presynaptic and intrinsic noise and showed that the balance of inhibitory and excitatory currents emerges at capacity. Zhang et al. (18) showed that many structural and dynamical properties of local cortical networks emerge in associative networks robustly loaded to capacity. These studies significantly differ from the present work both in terms of the models and the results. First, the model introduced in this article includes the complete error propagation cycle with presynaptic spiking errors combining with synaptic and intrinsic noises to produce errors in the neurons’ outputs. Second, the model allows for the possibility of having different levels of errors and noise during learning and memory retrieval. Third, the model makes it possible to analyze networks of neurons with heterogeneous properties. In terms of model results, we first show how errors and noise during learning facilitate reliable memory retrieval and next produce a comprehensive list of results related to network structure and dynamics that are then compared to the data from local cortical networks to validate the model and make predictions.

The manuscript is organized as follows. In Results, we first provide a description of an associative sequence learning model by a network of inhibitory and excitatory neurons in the presence of errors and noise in signal transmission and biologically inspired constraints on connectivity. Second, we solve the model numerically with nonlinear optimization for finite networks and analytically with the replica method in the thermodynamic limit and show how various sources of errors and noise affect the memory storage capacity of the neurons. Third, we describe the tradeoff between the capacity and reliability of stored memories and show that to retrieve the maximum amount of information memories must be loaded into the network in the presence of noise. Fourth, we examine the properties of connectivity and dynamics in model networks loaded to capacity with associative memory sequences in the presence of errors and noise and show that there is a region of parameters in which these properties agree with the reported experimental measurements. Fifth, we derive a perceptron-type learning rule that can be used to store associative memories into the network in an online manner and show that this biologically more plausible method leads to similar network properties as those obtained with the nonlinear optimization and replica methods. Finally, we examine the properties of networks of heterogeneous neurons and make predictions regarding network connectivity. The details of the replica calculation and numerical solutions of the associative memory storage model are provided in SI.

## RESULTS

### A. Network model of associative memory storage in the presence of errors and noise

We modeled associative sequence learning by a local (∼100 μm in size), all-to-all potentially (structurally) connected (29, 30) cortical network. The model network consisted of *N*_*inh*_ inhibitory and (*N* − *N*_*inh*_) excitatory McCulloch and Pitts neurons (17) (Figure 1A) and was faced with a task of learning a sequence of consecutive network states, *X*^1^ → *X*^2^ → …*X*^*m*+1^, in which *X*^*μ*^ is a binary vector representing target activities of all neurons at a time step *μ*, and the ratio *m*/*N* is referred to as the memory load. During learning, individual neurons had to independently learn to associate inputs they received from the network with the corresponding target outputs derived from the associative memory sequence. The neurons learned these input-output associations by adjusting the weights of their input connections, *J*_*ij*_ (weight of connection from neuron *j* to neuron *i*). In contrast to previous studies, we accounted for the fact that learning in the brain is accompanied by several sources of errors and noise. Within the model, these sources are divided into three categories (orange lightning signs in Figure 1A): (1) spiking errors, or errors in *X*^*μ*^, (2) synaptic noise, or noise in *J*_*ij*_, and (3) intrinsic noise, which combines all other sources of noise affecting the neurons’ postsynaptic potentials. The last category includes background synaptic activity and the stochasticity of ion channels. In the model, this category is equivalent to noise in the neurons’ thresholds of firing, *h*_*i*_ (for neuron *i*). In the following, asterisks are used to denote quantities containing errors or noise (e.g. *X*^\**μ*^), whereas symbols without asterisks represent the mean (for *h*_*i*_ and *J*_*ij*_) or target (for *X*^*μ*^) values.

The target neuron activities (e.g. binary scalar 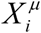) were independently drawn from neuron-dependent Bernoulli probability distributions: 0 with probability 1 – *f*_*i*_ and 1 with probability *f*_*i*_. Spiking errors in neuron activity states were introduced with the Bernoulli trials by making independent and random 1 to 0 changes with probabilities 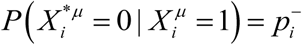 for spike failures and 0 to 1 changes with probabilities 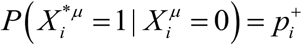 for erroneous spikes. Without loss of generality, we assumed that these two types of spiking errors are balanced, 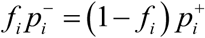, and do not affect the neuron’s firing probability, *f*_*i*_. This relation allowed us to describe both types of spiking errors in terms of the neuron’s overall spiking error probability, 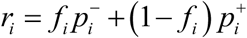:

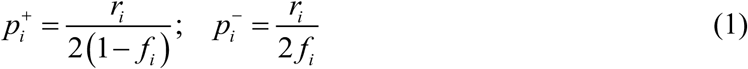

To describe synaptic noise, we followed the basic model of quantal synaptic transmission (31) and assumed that the variance of a given connection weight, 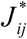, is proportional to its mean, 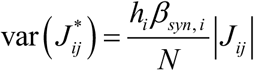. The dimensionless coefficient *β*_*syn, i*_ is referred to as the synaptic noise strength of neuron *i*, and the factor of *h*_*i*_ /*N* was introduced for convenience. We assumed that the intrinsic noise is Gaussian distributed across trials with the mean 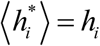 and variance 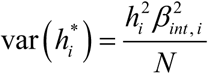. Here, *β*_*int, i*_ is a dimensionless coefficient called the intrinsic noise strength of neuron *i*, and, as before, a factor of 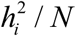 was introduced for convenience.

Similar to (27), two biologically inspired constraints were imposed on the learning process. First, the *l*_1_-norm of input connection weights of each neuron was fixed during learning, 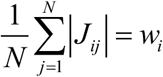. Here, parameter *w*_*i*_ is referred to as the average absolute connection weight of neuron *i*. Second, the signs of output connection weights of every neuron (inhibitory or excitatory) were fixed during learning, *J*_*ij*_ *g* _*j*_ ≥ 0. In these *N*^2^ inequalities, parameter *g* _*j*_ = 1 if neuron *j* is excitatory and –1 if it is inhibitory. Biological motivations for these constraints were previously discussed (27).

Individual neurons (e.g. neuron *i*) learned independently to associate noisy inputs they received from the network, *X*^\**μ*^, with the corresponding target outputs derived from the associative memory sequence, 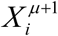. Neuron *i* is said to have learned the presented set of associations successfully if, in the presence of input spiking errors, synaptic and intrinsic noise, the fractions of its erroneous and failed spikes do not exceed its assigned spiking error probabilities, 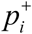 and 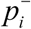 (Figure 1B). The above-described model for neuron *i* can be summarized as follows:

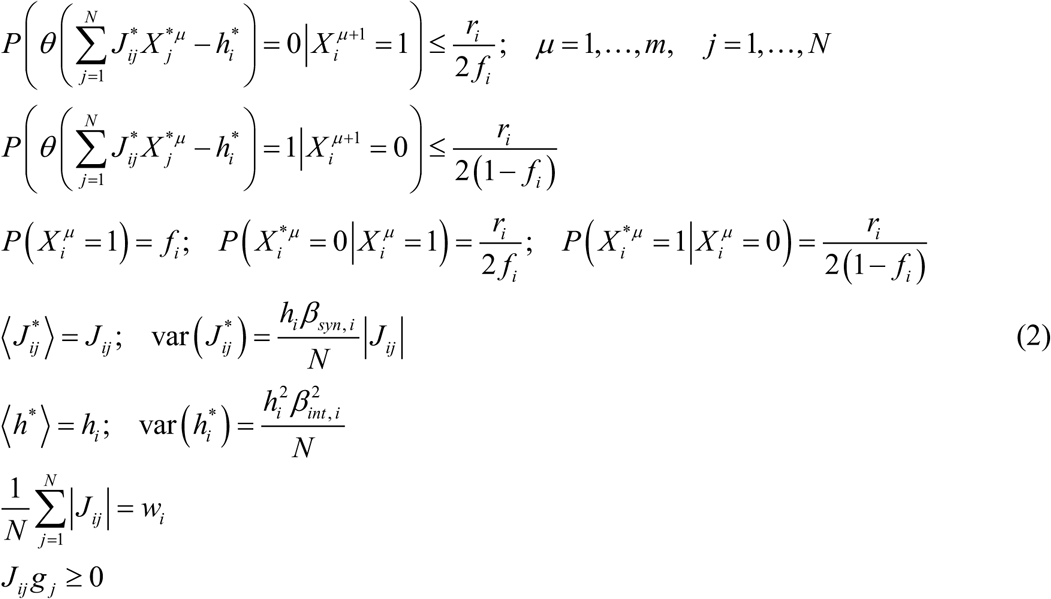

We note that, depending on the loaded associative memory sequence, Eqs. (2) may have multiple solutions if the learning problem faced by the neuron is feasible or no solution if the problem is not feasible. The neuron’s success probability in learning associative sequences of a given length is defined as the average of such binary outcomes over the sequences (Figure 1C). This probability is a decreasing function of the memory load and levels of errors and noise.

At the network level, the described associative memory storage model is governed by the network-related parameters *N* and {*g*_*i*_}, the memory load *m*/*N*, and the neuron-related parameters {*h*_*i*_}, {*w*_*i*_}, {*f*_*i*_}, {*r*_*i*_}, {*β*_*int, i*_}, and {*β*_*syn, i*_}, and the task is to find connection weights, {*J*_*ij*_}, that satisfy the requirements of Eqs. (2) for all neurons. In the following, we examine the properties of associative networks composed of inhibitory and excitatory neurons governed by the same and distributed sets of neuron-related parameters. We refer to these networks as homogeneous and heterogeneous.

### B. Solutions of the model

Eqs. (2) were solved with the replica method, nonlinear optimization, and perceptron-type learning rule (see SI). Each of these methods has its advantages and drawbacks, and, consequently, all three methods were used in this study. The replica method (32, 33) provides an analytical solution in the *N* → ∞ limit. Though neuron networks in the brain are finite, they are thought to be large enough to have many properties that are well described by this limit (18). More importantly, the analytical solution of the replica method reveals the dependence of the results on combinations of network parameters that can be explored with other methods. The downside of the replica solution is that it does not provide the full connectivity matrix, *J*_*ij*_, but instead gives the connectivity statistics that cannot be readily used to calculate all relevant network properties. Nonlinear optimization can be used to solve Eqs. (2). This method is fast and accurate for small networks, yielding the full connectivity matrix, but is impractical for large networks (*N* ∼ 1,000). As the replica and nonlinear optimization solutions cannot be readily implemented by neural networks in the brain, we also developed a biologically more realistic perceptron-type learning rule that can be used to approximate the solution of Eqs. (2). Because simulations based on the perceptron-type learning rule are time-consuming, results for varying levels of errors and noise were obtained with the replica and nonlinear optimization methods, while the perceptron-type learning rule was used only for a biologically realistic set of parameters in order to confirm that all three methods lead to similar results.

In the *N* → ∞ limit, the associative memory storage problem for a neuron loaded to capacity was solved with the replica method (see sections E-G of SI). This solution for a neuron in a homogeneous network (*f*_*i*_ = *f, r*_*i*_ = *r, β*_*syn, i*_ = *β*_*syn*_, and *β*_*int, i*_ = *β*_*int*_) depends on the following combination of the intrinsic and synaptic noise strengths (see section H of SI):

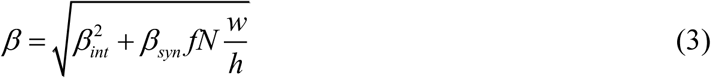

The latter is referred to as the postsynaptic noise strength. In the following, we assume that the postsynaptic noise strength, *β*, and the spiking error probability, *r*, can differ between the times of learning and memory retrieval and add subscripts “*learn*” and “*retr*” to these parameters to distinguish among the two phases.

Figure 1C shows that when the memory load is relatively low, the probability of successful learning by a neuron is close to 1. With increasing load, the learning problem becomes more difficult, and the success probability undergoes a smooth transition from 1 to 0. Memory load corresponding to the success probability of 0.5 is referred to as the neuron’s associative memory storage capacity, *α*. This capacity depends on the levels of errors and noise accompanying learning and other parameters of the model. With increasing network size, *N*, the transition from successful learning to inability to accurately learn the complete memory sequence becomes sharper, and the neuron’s capacity monotonically approaches its *N* → ∞ limit, which is referred to as the critical capacity, *α*_*c*_. Figure 1D illustrates the dependence of a single-neuron critical capacity on the postsynaptic noise strength and spiking error probability. As expected, because noise makes the learning problem more challenging, *α*_*c*_ is a decreasing function of *β*_*learn*_. Dependence of *α*_*c*_ on *r*_*learn*_ is more complex as this parameter controls both the presynaptic and output spiking errors of the neuron. On the one hand, an increase in *r*_*learn*_ makes the learning problem more challenging as the presynaptic errors increase and on the other, the problem becomes easier due to a higher tolerance of output errors.

### C. The tradeoff between capacity and reliability of loaded memories

Can memories loaded into individual neurons be successfully recalled at the network level? To answer this question, we loaded neurons in the network to capacity with associations derived from a single associative sequence by solving Eqs. (2). The postsynaptic noise and spiking errors during learning were set at the levels *β*_*learn*_ and *r*_*learn*_ described by the green asterisk in Figure 1D. During memory retrieval, the network was initialized at the beginning of the loaded sequence, *X*^1^, and no additional spiking errors, beyond those produced by the network at subsequent steps, were added during memory playout. At each step of memory playout, synaptic and intrinsic noise were added independently to every connection and every neuron in the network at strengths governed by *β*_*retr*_, which can differ from *β*_*learn.*_

The sequence is said to be retrieved completely if the network states during the retrieval do not deviate substantially from the target states. Otherwise, the sequence is said to be retrieved partially, and the retrieved sequence length is defined by the number of steps taken to the point where the network states begin to deviate substantially from the target states (Figure 2A). In practice, there is no need to precisely define the threshold amount of deviation. This is because for large networks the fraction of errors in a retrieved network state either fluctuates around 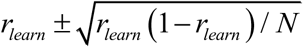 (mean ± SD) or diverges to 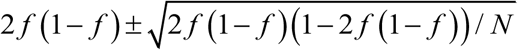 (expected fraction of differences in two random network states of firing probability *f*), which is significantly greater for the chosen values of parameters *r*_*learn*_ and *f*. Figure 2B shows the probability of retrieving the complete loaded sequence and the fraction of retrieved sequence length for different values of *β*_*learn*_. It illustrates that memory sequences can be reliably retrieved if they were loaded with the postsynaptic noise strength that is slightly higher than that during memory retrieval. Likewise, the averaged retrieved sequence length fraction increases with *β*_*learn*_ and approaches one as the latter exceeds the noise strength present during retrieval. A similar conclusion can be drawn from Figure 2C which shows the map of the retrieval probability as a function of *β*_*learn*_ and *r*_*learn*_. Errors and noise during learning make memory retrieval more reliable. However, the reliability of loaded memories comes at the expense of the memory storage capacity, *α*. Figure 2D shows the tradeoff between the retrieval probability and capacity of loaded associative memories in which higher levels of errors and noise during learning enable reliable memory retrieval but reduce *α*.

**Figure 2:**
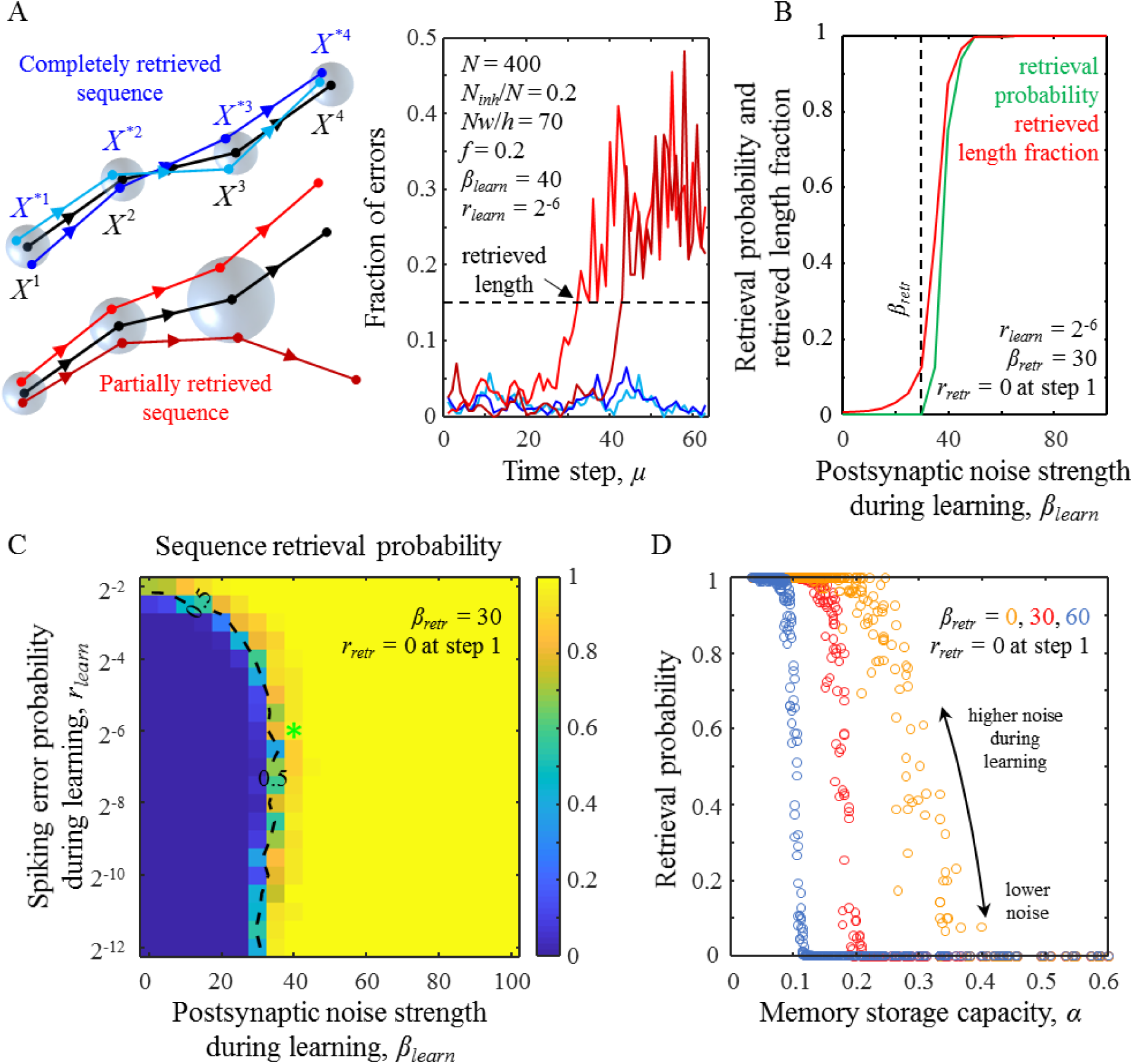
Retrieval of loaded associative memory sequences and the tradeoff between capacity and reliability of loaded memories. **A.** Illustration of memory playout during complete and partial memory retrieval (left). The target memory sequence is shown in black, while the sequences retrieved on different trials are in blue and red. Memory retrieval is incomplete when the retrieved sequence deviates significantly from the target sequence (see text for details). Radii of blue spheres illustrate the root-mean-square Euclidean distances between the retrieved and target states. The fraction of errors as a function of time step during sequence retrieval (right). Successfully retrieved sequences do not deviate from the loaded sequences by more than a threshold amount (dashed line). The parameters of the associative network are provided in the figure. The values of *β*_*learn*_ and *r*_*learn*_ correspond to the green asterisk from Figure 1D. **B.** The probability of successful memory retrieval (green) and the retrieved fraction of loaded sequence length (red) as a function of *β*_*learn*_. The postsynaptic noise strength *β*_*retr*_ = 30 (dashed line) at every step of memory retrieval and *r*_*retr*_ was set to 0 at the first step. **C**. Map of retrieval probability as a function of *β*_*learn*_ and *r*_*learn*_. Dashed isocontour is drawn as a guide to the eye. The location of the green asterisk is the same as in Figure 1D. **D.** The tradeoff between memory retrieval probability and *α*. Individual points correspond to all values of *β*_*learn*_ and *r*_*learn*_ considered in (C). Higher errors and noise during learning result in lower *α* and higher retrieval probability regardless of the noise strength during memory retrieval (different colors). The results shown in (A-D) were obtained with the nonlinear optimization method described in section I of SI. For every parameter setting, the results shown in (B-D) were averaged over 100 networks and 1,000 retrievals of the loaded sequence in each network.

### D. Noise during learning is required for optimal retrieval of stored information

Figures 3A, B show the maps of expected retrieved information per sequence playout calculated in two different ways. In the first calculation, the contribution of partially retrieved sequences to the expected retrieved information was set to zero, while in the second, partially retrieved sequences contributed in proportion of the retrieved sequence length (see section K of SI for details). Both maps illustrate that optimal retrieval of stored information is achieved when memories are stored in the presence of noise, *β*_*learn*_ > 0. This conclusion is independent of the postsynaptic noise strength during memory retrieval, which was set to *β*_*retr*_ = 30 in Figures 3A, B. To illustrate this finding, we averaged the maps over the *r*_*learn*_ dimension and determined *β*_*learn*_ that correspond to the maxima of the retrieved information. Figure 3C illustrates the results of this procedure for different values of *β*_*retr*_, showing that the optimal *β*_*learn*_ is greater than zero even when there is no noise during memory retrieval. The optimal *β*_*learn*_ increases with *β*_*retr*_, and when noise is high the two noise strengths approximately equal.

**Figure 3:**
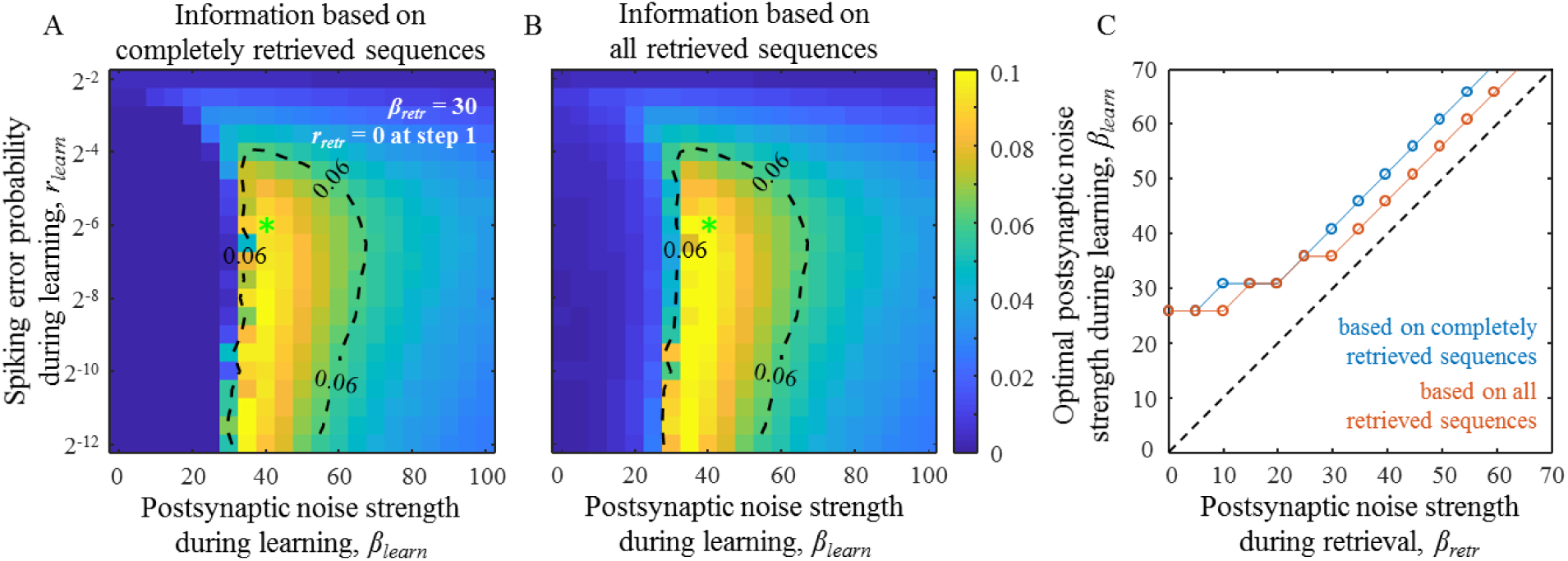
Postsynaptic noise during learning is required for optimal retrieval of stored of information. **A-B.** Maps of expected retrieved information per memory playout calculated based on completely retrieved sequences (A) and completely and partially retrieved sequence (B) in bits×*N*^2^ as functions of *β*_*learn*_ and *r*_*learn*_. *β*_*retr*_ = 30 at every step of memory retrieval and *r*_*retr*_ was set to 0 at the first step. Dashed isocontours are drawn as guides to the eye. The locations of the green asterisks are the same as in Figure 1D. **C.** The maximum of retrieved information is achieved when *β*_*learn*_ is greater than zero regardless of the value of *β*_*retr*_. The optimal postsynaptic noise strengths were calculated based on the averages of the results from (A) (blue line) and (B) (orange line) over the range of *r*_*learn*_ values from (A, B). All results were obtained with the nonlinear optimization method described in section I of SI and averaged over 100 networks and 1,000 retrievals of the loaded sequence in each network for every parameter setting.

### E. Neuron-to-neuron connectivity in associative networks of homogeneous inhibitory and excitatory neurons

One of the most salient features of sign-constrained associative learning models, such as the one described in this study, is that finite fractions of inhibitory and excitatory connections assume zero weights at capacity (25), mirroring the trend observed in many local cortical networks. We compared the connection probabilities (*P*_*con*_) and the coefficients of variation (CV) of non-zero connection weights in associative networks at capacity to the connection probabilities and CVs of unitary postsynaptic potentials (uPSP) obtained experimentally. To that end, we used the dataset compiled in (18) based on 87 electrophysiological studies describing neuron-to-neuron connectivity for 420 local cortical projections (lateral distance between neurons < 100 μm). Figure 4A shows that the average inhibitory *P*_*con*_ (38 studies, 9,522 connections tested) is significantly larger (p < 10^−10^, two-sample t-test) than the average excitatory *P*_*con*_ (67 studies, 63,020 connections tested). Associative networks exhibit a similar trend in the entire region of considered *β*_*learn*_ and *r*_*learn*_ values (Figures 4B, C). What is more, in the (*β*_*learn*_, *r*_*learn*_) parameter region demarcated with the dashed isocontours and arrows in Figures 4B, C, the model results are consistent with the middle 50% of the experimentally measured *P*_*con*_ values for inhibitory and excitatory connections.

**Figure 4:**
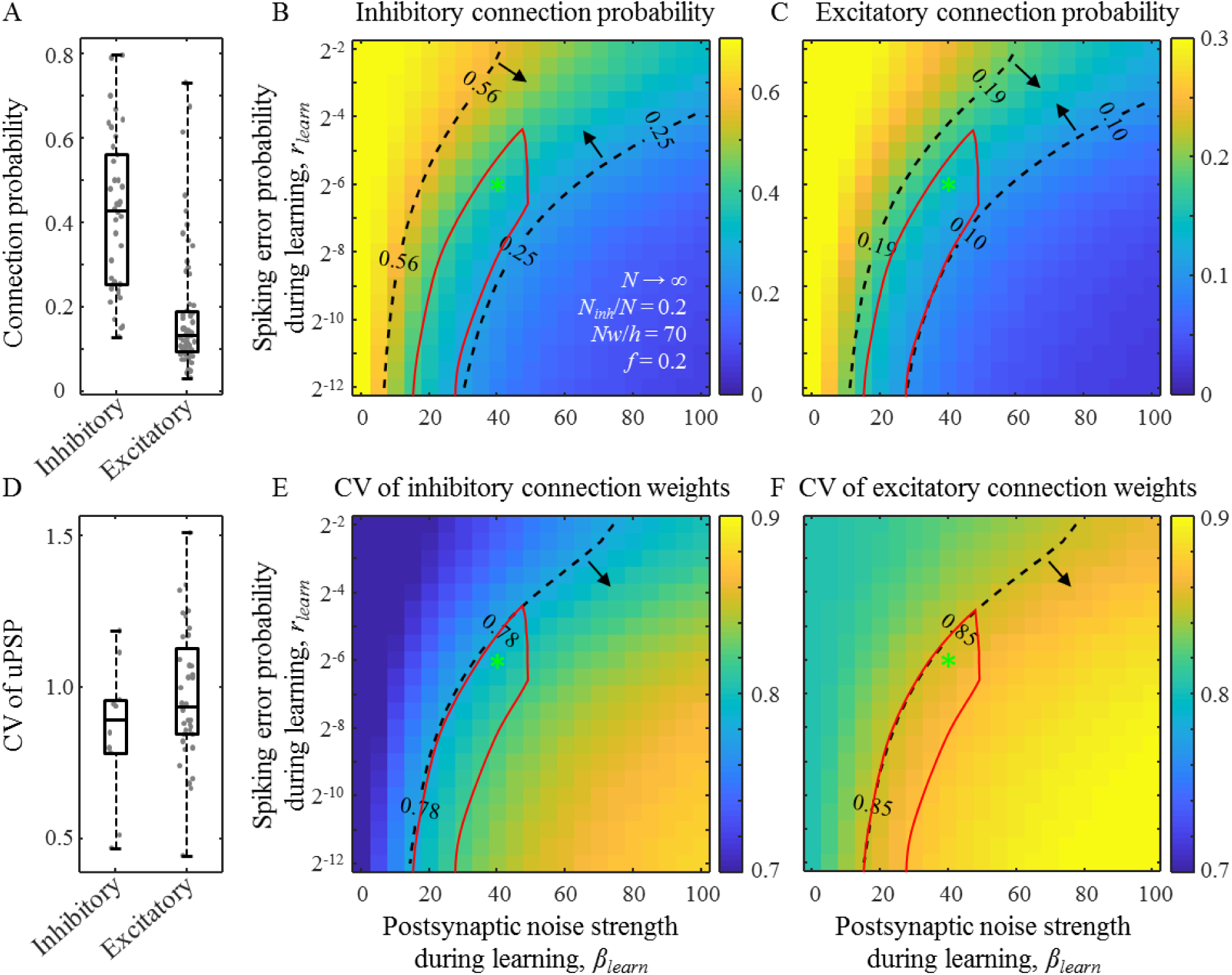
Comparison of structural properties of the model and cortical networks. **A.** Inhibitory and excitatory connection probabilities reported in 87 studies describing 420 local cortical projections. Each dot represents the result of a single study/projection. **B, C.** Maps of inhibitory and excitatory connection probabilities as functions of *β*_*learn*_ and *r*_*learn*_. The results are based on the replica method (see section H of SI). Dashed isocontours and arrows illustrate the interquartile ranges of the experimentally observed connection probabilities from (A). The red contour outlines a region of parameters that is consistent with all structural and dynamical measurements in cortical networks considered in this study. The locations of the green asterisks are the same as in Figure 1D. **D**-**F.** Same for the CV of inhibitory and excitatory connection weights. (A) and (D) were adapted from (18).

Figure 4D shows that the average CV of inhibitory uPSP (10 studies, 503 connections recorded) is slightly lower than that for excitatory (36 studies, 3,956 connections recorded), and this trend is also reproduced by the associative networks in the entire region of considered *β*_*learn*_ and *r*_*learn*_ values (Figures 4E, F). As before, there are (*β*_*learn*_, *r*_*learn*_) parameter regions in these maps in which the results of the model are consistent with the middle 50% of the CV of uPSP measurements for inhibitory and excitatory connections.

### F. Spontaneous dynamics in associative networks of homogeneous inhibitory and excitatory neurons

The model associative networks can exhibit irregular and asynchronous spiking activity like that observed in cortical networks. To analyze such spontaneous (not learned) network dynamics, we used associative networks loaded to capacity, initialized them at random states of firing probability *f* = 0.2, and followed their activity for 1,000 time-steps. Because the number of available network states, which is exponential in *N*, is much larger than the number of loaded states, *αN*, the spontaneous network activity in the numerical simulations never passed through any of the loaded states.

To quantify the degree of similarity in the dynamics of the model and brain networks we compared the CV of inter-spike-intervals (ISI) and the cross-correlation coefficient of spiking neuron activity in the model to those measurements obtained experimentally. Dashed isocontour in Figure 5A outlines (*β*_*learn*_, *r*_*learn*_) parameter region in which the model CV of ISI is consistent with the 0.7-1.1 range measured in different cortical systems (34-38). Similarly, Figure 5B shows that there is a (*β*_*learn*_, *r*_*learn*_) parameter region in which the calculated spike cross-correlation coefficients are in agreement with the interquartile range of the corresponding cortical measurements, 0.04-0.15 (39). The degree of asynchrony in spontaneous spiking activity in associative networks increases with the postsynaptic noise strength, which can be explained by the decrease in connection probability (Figures 4B, C) and, consequently, a reduction in the amount of common input to the neurons.

**Figure 5:**
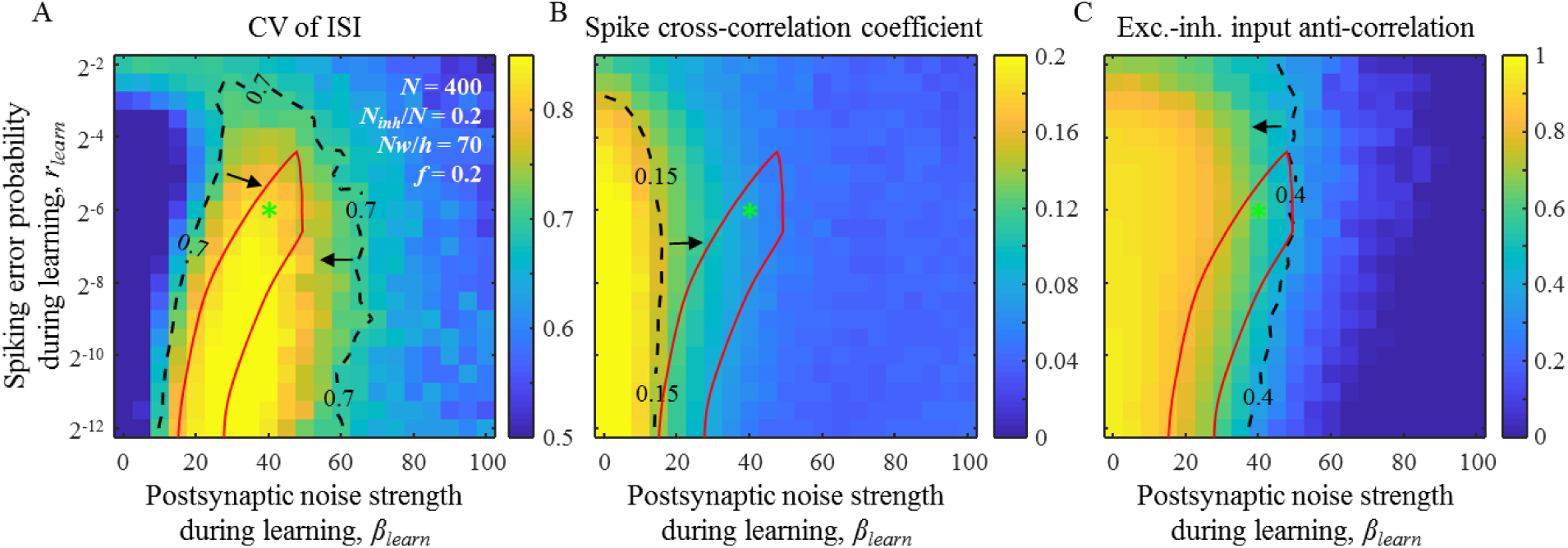
Comparison of dynamical properties of the model and cortical networks. **A.** The CV of ISI for spontaneous (not learned) activity as a function of *β*_*learn*_ and *r*_*learn*_. Results were obtained with nonlinear optimization (see section I of SI). Dashed isocontour and arrows demarcate a region of CV values that is in general agreement with experimental measurements. **B.** Same for the cross-correlation coefficient of neuron spike trains. **C.** Same for the anti-correlation coefficient of inhibitory and excitatory postsynaptic inputs to a neuron. The red contour outlines a region of parameters that is consistent with the considered structural and dynamical measurements. The locations of the green asterisk are the same as in Figure 1D. All results were obtained with the nonlinear optimization method described in section I of SI and averaged over 100 networks and 100 runs for each network for every parameter setting.

It was shown that irregular and asynchronous activity can result from the balance of inhibitory and excitatory postsynaptic inputs to individual cells (40, 41). In a balanced state, the magnitudes of these inputs are much greater than the threshold of firing, but, due to a high degree of anti-correlation, these inputs nearly cancel, and firing is driven by fluctuations. Figure 5C shows a region of parameters in which neurons in the associative model function in a balanced regime. Because it is difficult to simultaneously measure inhibitory and excitatory postsynaptic inputs to a neuron, the anti-correlation of inhibitory and excitatory inputs has only been measured in nearby cells, averaging to about 0.4 (42, 43). As within-cell anti-correlations are expected to be stronger than between-cell anti-correlations, 0.4 was used as a lower bound for the former (dashed isocontour and arrow in Figure 5C).

The seven error-noise regions obtained based on the properties of neuron-to-neuron connectivity (Figure 4) and network dynamics (Figure 5) have a non-empty intersection (red contour in Figures 4 and 5). In this biologically plausible region of parameters, the considered properties of the associative networks are consistent with the corresponding experimental measurements. This observation suggests that *β*_*learn*_ must lie in the 20-50 range and *r*_*learn*_ must be less than 0.06. While we are not aware of direct experimental measurements of these parameters, the low value of *r*_*learn*_ is in qualitative agreement with the reliability of firing patterns evoked by time-varying stimuli *in vivo* (38) and *in vitro* (44).

### G. Solution of the model with a perceptron-type learning rule

As the replica and nonlinear optimization solutions of Eqs. (2) cannot be easily implemented by neural networks in the brain, we set out to develop a biologically more plausible online solution to the associative learning problem. The following perceptron-type learning rule was devised to approximate the solution of Eqs. (2) (see section J of SI). At each learning step, e.g. *μ*, a neuron receives an input containing spiking errors, *X*^\**μ*^, combines it with postsynaptic noise, and produces an output, *y*^\**μ*^. If this output differs from the neuron’s target output, *y*^*μ*^, the neuron’s input connection weights are updated in four consecutive steps:

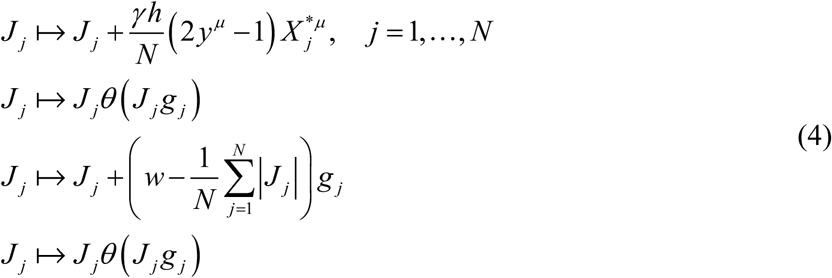

The first line in Eqs. (4) is a stochastic perceptron learning step (45), in which parameter *γ* is referred to as the learning rate. The second line enforces the sign constraints, while the last two lines combined implement the homeostatic *l*_1_-norm constraint and are equivalent to the soft thresholding used in LASSO regression (46). In contrast to the standard perceptron learning rule, Eqs. (4) utilize noisy inputs and enforce sign and homeostatic constraints at every learning step.

By including presynaptic spiking errors, synaptic and intrinsic noise in the condition that triggers the learning steps outlined in Eqs. (4), the learning rule implicitly depends on the model parameters {*r*_*j*_}, {*β*_*syn, j*_} describing the fluctuations in the neuron’s inputs (indexed with *j*), and the parameter *β*_*int*_ which describes the neuron’s intrinsic noise. Because Eqs. (4) are designed to approximately minimize the neuron’s output spiking error probability for a given memory load (see section J of SI), which at capacity matches the desired output error probability of the neuron *r*, the learning rule also depends implicitly on fluctuations in the neuron’s output.

Figure 6 compares the theoretical solution obtained with the replica method in the *N* → ∞ limit with numerical solutions for networks of *N* = 200, 400, and 800 neurons obtained with nonlinear optimization and the perceptron-type learning rule. Figure 6A shows that the perceptron-type learning rule sometimes fails to find a solution to a feasible learning problem, i.e. a problem that can be solved with nonlinear optimization. Yet, even in such cases, the perceptron connection weights in a steady state (after 10^6^ learning epochs) are well-correlated with the nonlinear optimization weights (Figure 6B). Therefore, though the perceptron-type learning rule is not as efficient as nonlinear optimization, it can find an approximate solution to the learning problem. Consistent with this conclusion, the associative memory storage capacity of a neuron loaded with the perceptron-type learning rule is 15% - 18% lower than that loaded with nonlinear optimization, and the two methods lead to similar structural and dynamical network properties (red and blue bars in Figure 6C).

**Figure 6:**
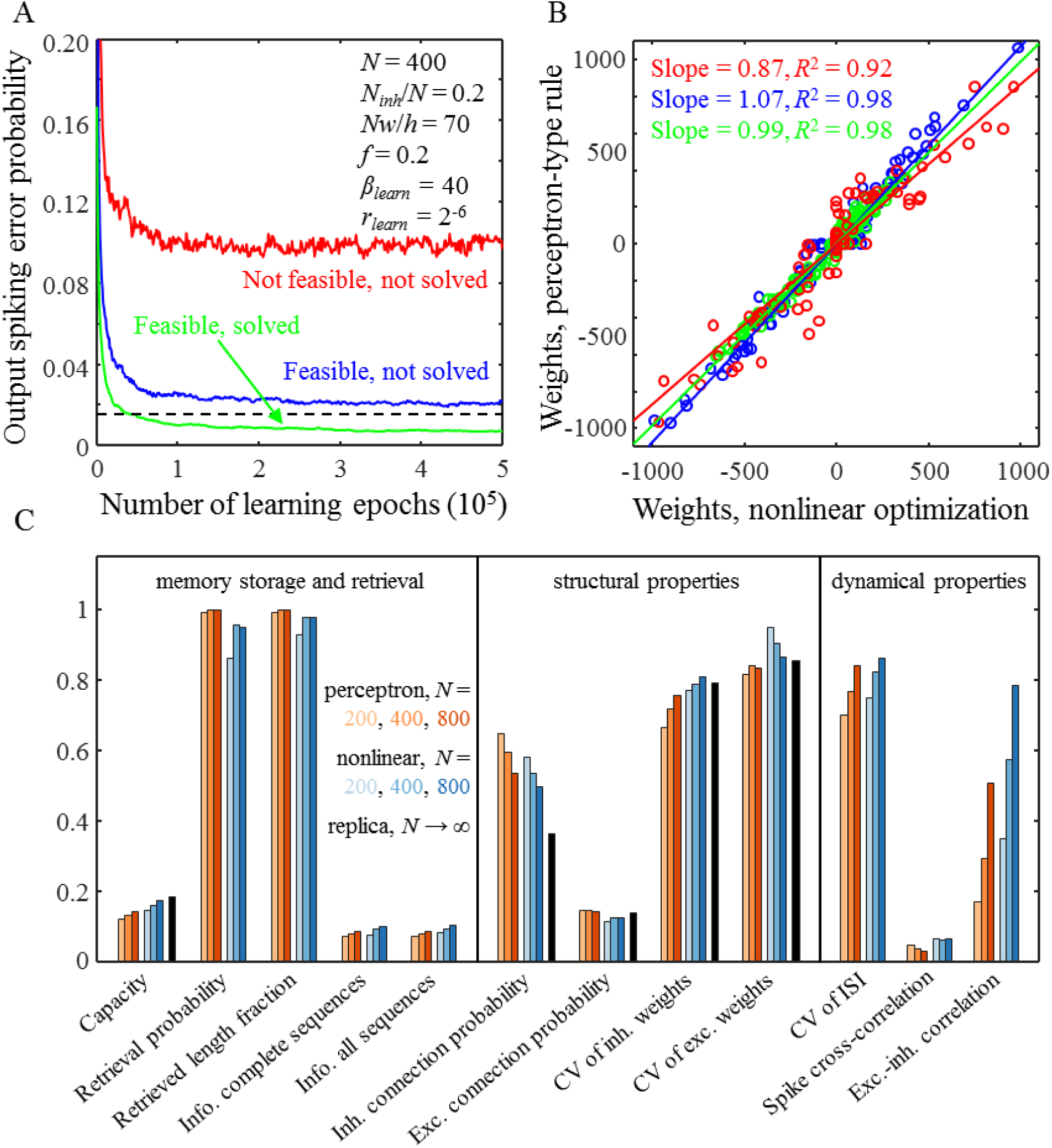
Comparison of solutions obtained with the perceptron-type learning rule, nonlinear optimization, and replica method. **A.** Output error probability as a function of the number of learning epochs for the perceptron-type learning rule. The black dashed line indicates the target output error probability. Results for three different cases are shown: a not feasible problem (red line), a feasible problem which was not solved with the perceptron-type learning rule (blue line), and a feasible problem which was solved with the perceptron-type learning rule (green line). The parameters of the associative network are provided in the figure. The values of *β*_*learn*_ and *r*_*learn*_ correspond to the green asterisk from Figure 1D. **B.** Comparisons of connection weights obtained with the perceptron-type learning rule and nonlinear optimization for the three cases shown in (A). Straight lines are the best linear fits. **C.** Comparisons of memory storage capacity, retrieval, structural, and dynamical properties in networks of *N* = 200, 400, and 800 neurons obtained with the perceptron-type learning rule (red colors) and nonlinear optimization (blue colors). The memory storage capacity and structural properties calculated with the replica method in the *N* → ∞ limit are shown in black.

### H. Properties of heterogeneous associative networks

The associative learning model, Eqs. (2), makes it possible to investigate the properties of networks composed of heterogeneous populations of inhibitory and excitatory neurons. Specifically, we examined the effects of distributed spiking error probabilities and distributed synaptic and intrinsic noise strengths on properties of connectivity at critical capacity. Figures 7A-C show that in networks of neurons with heterogeneous spiking error probabilities (homogeneous in all other parameters), the probabilities and weights of inhibitory and excitatory connections monotonically decrease with increasing *r*_*learn*_. Therefore, as may have been expected, connections originating from unreliable presynaptic neurons (high *r*_*learn*_) are depressed and/or eliminated during learning. Properties of networks of neurons with distributed synaptic and intrinsic noise strengths (homogeneous otherwise) depend on the combination of these parameters in the form of the postsynaptic noise strengths, *β*_*learn*_. Figures 7D-F show how connection probabilities and average connection weights depend on *β*_*learn*_. Similar to the previous case, connections onto noisy neurons (high *β*_*learn*_) are less probable. Here, however, the average inhibitory and excitatory connection weights increase with *β*_*learn*_ due to the homeostatic *l*_1_-norm constraint, Eqs. (2).

**Figure 7:**
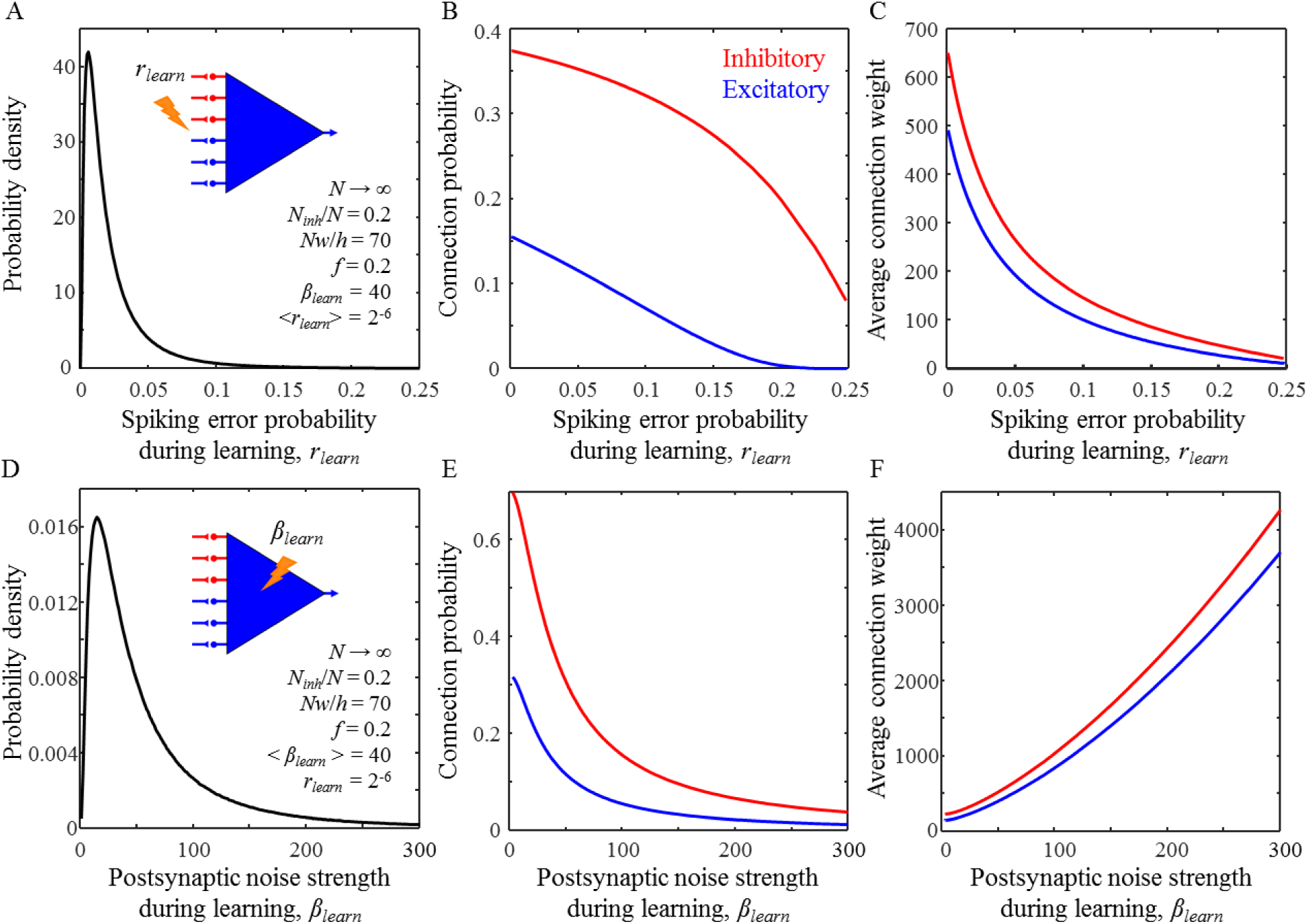
Properties of connections in associative networks of heterogeneous neurons. **A-C.** Connection probability (B) and average non-zero connection weight (C) for inhibitory (red) and excitatory (blue) connections in a network of neurons with distributed spiking error probabilities and homogeneous in all other parameters. The spiking error probabilities of inhibitory and excitatory inputs during learning were randomly drawn from the log-normal distribution shown in (A). Unreliable inputs have lower probabilities and weights. The parameters of the associative network are shown in (A). The values of *β*_*learn*_ and *<r*_*learn*_> correspond to the green asterisk from Figure 1D. **D-F.** Same for a network of neurons with heterogeneous postsynaptic noise strengths. The postsynaptic noise strengths of neurons during learning were randomly drawn from the log-normal distribution shown in (D). Noisy neurons receive stronger but fewer inhibitory and excitatory inputs.

Motivated by the agreement between the results of the associative learning model and cortical measurements, we put forward two predictions that can be tested in future experiments. First, we predict that in cortical networks, inhibitory and excitatory connections originating from unreliable neurons or neuron classes must have lower connection probabilities and average uPSPs (Figures 7B, C). Second, we predict that connections onto noisy neurons or neuron classes must have lower connection probabilities but higher average uPSPs (Figures 7E, F).

## DISCUSSION

This study incorporates a comprehensive description of the error propagation cycle into the model of associate sequence learning by recurrent networks of neurons with biologically inspired constraints. It shows that errors and noise during learning can be beneficial, as they can increase the reliability of loaded memories to fluctuations during memory retrieval. Because errors and noise are both free and unavoidable harnessing their power, rather than trying to suppress it, may be an efficient way of improving the reliability of memories in the brain. This mechanism is illustrated in Figure 8. When the associative memories are loaded at a below capacity level, the solution region of Eqs. (2) is relatively large. A solution, e.g. a vector of connection weights of a neuron obtained with a perceptron-type learning rule, may be located near the solution region boundary. Such a solution is deemed unreliable because a small amount of noise during memory retrieval can move it outside the solution region, resulting in spiking errors that can disrupt the associative sequence retrieval process (Figures 8A). By adding noise during learning, the solution can be forced to move away from the boundary, thus making it more reliable (Figures 8B). However, increasing the noise strength reduces the neuron’s capacity, and at a certain strength, the capacity and memory load are guaranteed to match (Figure 8C). A further increase in noise strength can improve the reliability even more, but at the expense of the memory load as the latter must remain at or below capacity (Figure 8D). An alternative way of improving reliability is by suppressing noise during memory retrieval (Figure 8E). Interestingly, it has been shown that visual attention that improves behavioral performance reduces the variability in spike counts of individual neurons in Macaque V4 (5, 6). Though significant, the amount of reduction is relatively small, suggesting that this mechanism has physical limitations. Therefore, using noise during learning can enhance the reliability of stored memories beyond what can be accomplished by attending to the memory retrieval process.

**Figure 8:**
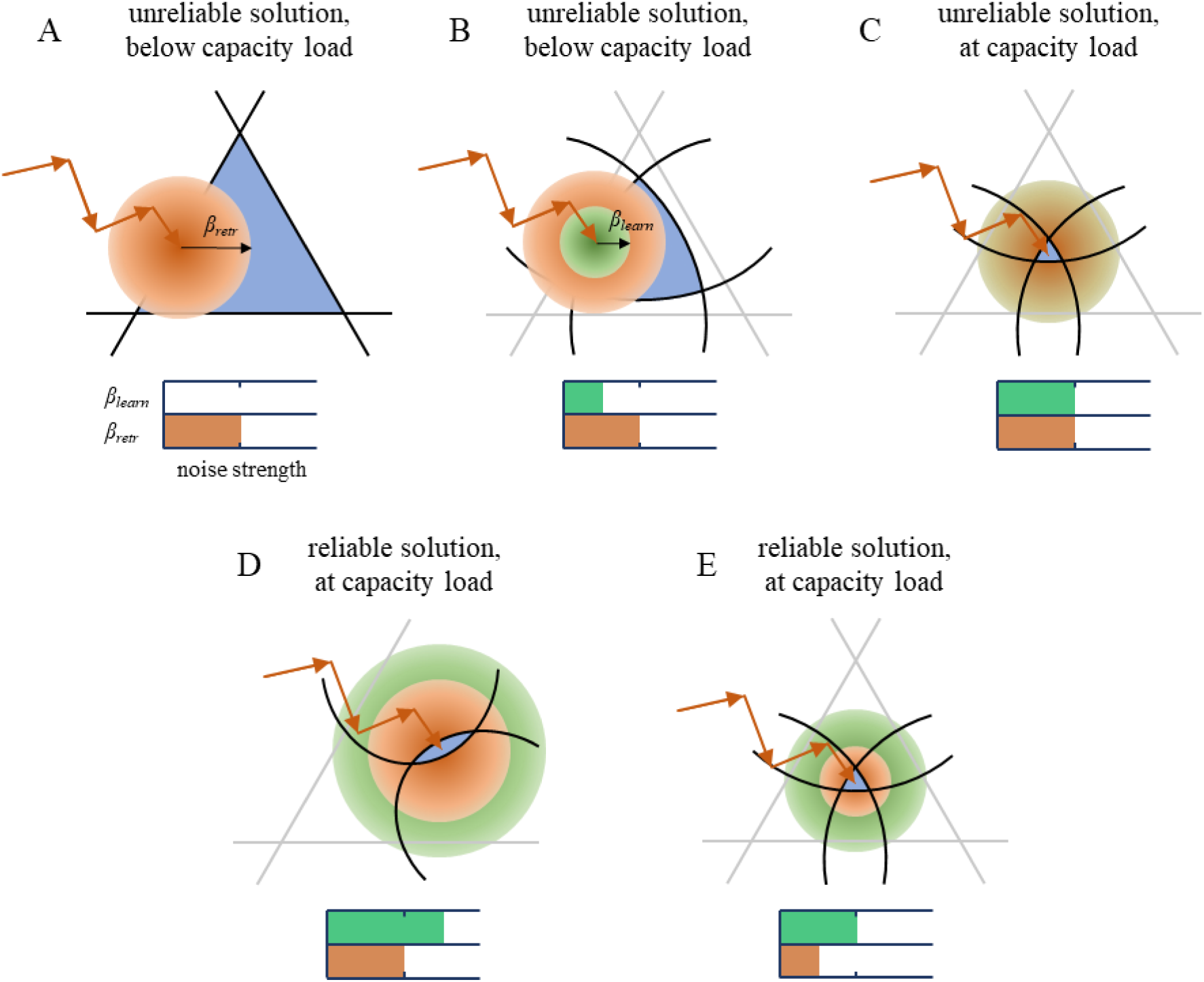
Increasing the noise strength during learning or decreasing it during memory recall leads to more reliable solutions. **A.** The associative learning problem for a below capacity load in the absence of noise during learning, *β*_*learn*_ = 0. The solution region (blue) is bounded by hyperplanes corresponding to the individual associations (black lines). The learning phase (red arrows) ends as the connection weight vector enters the solution region. The solution shown in (A) is unreliable because noise during memory retrieval (red cloud of radius *β*_*retr*_) can move it outside the solution region with high probability. **B.** Adding noise during learning (green cloud of radius *β*_*learn*_) transforms the association hyperplanes (gray lines) into hypersurfaces (black lines), Eqs. (2), reducing the solution region and forcing the connection weight vector further away from the hyperplanes. This increases solution reliability. **C.** The continued increase of the noise strength improves reliability as the solution region shrinks to zero. At this noise strength, the memory load is at capacity. A further increase in reliability can be achieved by increasing the noise strength during learning (D) or decreasing it during retrieval (E). In the former case, the memory load must be reduced to match the reduction in capacity.

The model described in this study assumes that individual neurons learn independently from one another and are loaded with memories to capacity. There is no direct support for these assumptions, but they have been shown to lead to structural and dynamical network properties that are consistent with experimental data (9, 18, 26, 28). This study corroborates these assumptions by matching a variety of experimental results with a single set of model parameters. The derived perceptron-type rule can mediate learning by modifying connection weights based on local activities of pre- and postsynaptic neurons in the presence of errors and noise. This is biologically feasible; however, a supervision signal must be fed to each neuron during learning. This is a major drawback of the presented rule, and the supervised learning rules in general because the origins of this signal in the brain remain unknown.

## ACKNOWLEDGMENTS

This work was supported by the AFOSR grant FA9550-15-1-0398 and the NSF grant IIS-1526642.

## SUPPORTING INFORMATION

This Supporting Information describes a model of associative memory storage by a McCulloch and Pitts neuron (17) receiving inhibitory and excitatory inputs (Figure 1A, bottom). In contrast to previous studies [see e.g. (9, 10, 18, 22, 23, 26-28)], the model explicitly accounts for several distinct sources of errors and noise present during learning and incorporates two biologically inspired constraints on connectivity. The model was solved theoretically with the replica method in the limit of infinite network size (32, 33) and numerically, with nonlinear optimization and perceptron-type learning rule, for large but finite networks.

### A. Perceptron model with biologically inspired constraints in the presence of errors and noise

We consider a single perceptron-like neuron that receives *N*_*inh*_ inhibitory and (*N − N*_*inh*_) excitatory input connections (Figure 1A). The neuron is faced with a task of learning a set of *m* input-output associations {*X*^*μ*^, *y*^*μ*^}, where *X*^*μ*^ and *y*^*μ*^ are a binary vector and a scalar describing the neuron’s input and target output at time step *μ*. The neuron must learn *m* input-output associations by adjusting the weights of its connections in the presence of two constraints. First, the *l*_1_-norm of connection weights must remain fixed during learning. Second, the signs of inhibitory and excitatory connection weights must not change during learning. In the following, *J*_*j*_ denotes the weight of connection *j*.

Learning in the model is accompanied by four types of errors and noise. These include presynaptic and output spiking errors, or errors in *X*^*μ*^ and *y*^*μ*^, synaptic noise, or noise in *J*, and intrinsic noise, or noise in the neuron’s threshold of firing, *h*. In the following, we use asterisks to denote quantities containing errors or noise (e.g. *X*^\**μ*^), whereas variables without asterisks represent the mean (for *h* and *J*_*j*_) or target (for *X*^*μ*^ and *y*^*μ*^) values.

Individual elements of 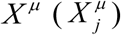, are independently drawn from connection-dependent Bernoulli probability distributions: 0 with probability 1 – *f*_*j*_ and 1 with probability *f*_*j*_. Errors in *X*^*μ*^ are introduced with Bernoulli trials by making independent and random changes in 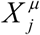 with probabilities 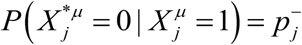 and 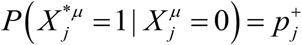. Similarly, the target outputs, *y*^*μ*^, are independently drawn from a Bernoulli probability distribution: 0 with probability 1 – *f*_*out*_ and 1 with probability *f*_*out*_. The probabilities of error in the neuron’s output are bounded, 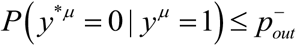 and 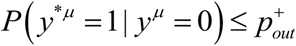 (Figure 1B). In the following, we only consider the output spiking error probabilities in the ranges 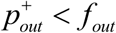 and 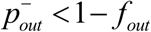, which is required for the stability of the replica solution. Noises in the neuron’s connection weights and postsynaptic potential are described in the main text. The resulting model is equivalent to Eqs. (2) with minor changes in the notation: 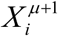 is replaced with *y*^*μ*^, *f*_*i*_ with *f*_*out*_, 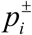 with 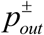, and index *i* is dropped. The model can be summarized as follows:

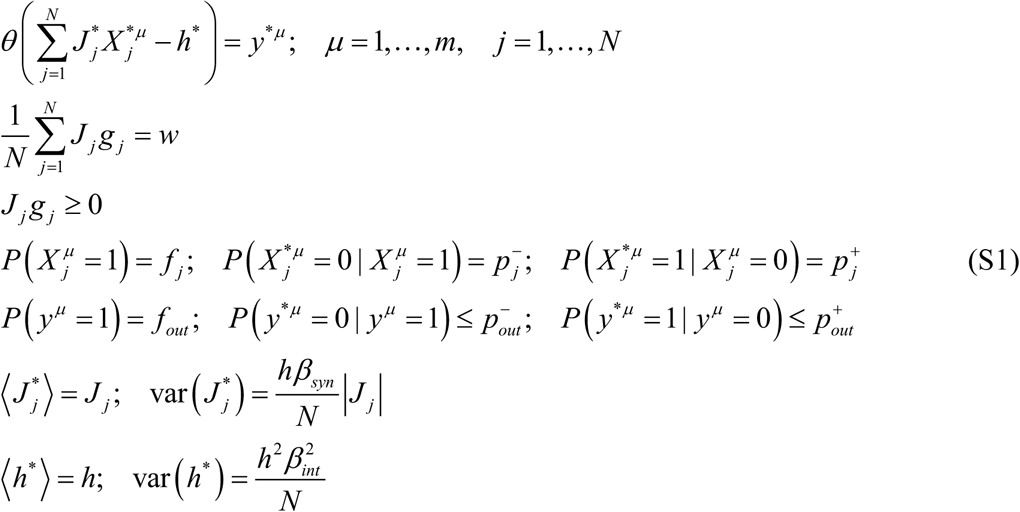

In these expressions, *θ* is the Heaviside step-function. To enforce sign-constraints on connection weights we introduced scalars *g*_*j*_, equaling 1 if the input *j* is excitatory and –1 if it is inhibitory. Parameter *w* is referred to as the average absolute connection weight. The neuron is faced with the task of finding the connection weights, {*J* _*j*_}, that satisfy Eqs. (S1) for a given set of model parameters: 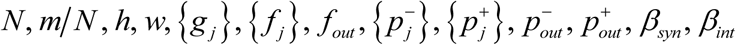.

### B. Reformulation of the model in the large *N* limit

In the limit of large *N*, the Central Limit Theorem ensures that the neuron’s postsynaptic potential (PSP), 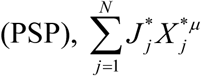, is Gaussian distributed at every time step. Therefore, the deviation of PSP from the threshold of firing, 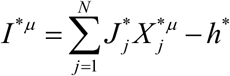, is also Gaussian distributed with the mean and standard deviation given by the following expressions:

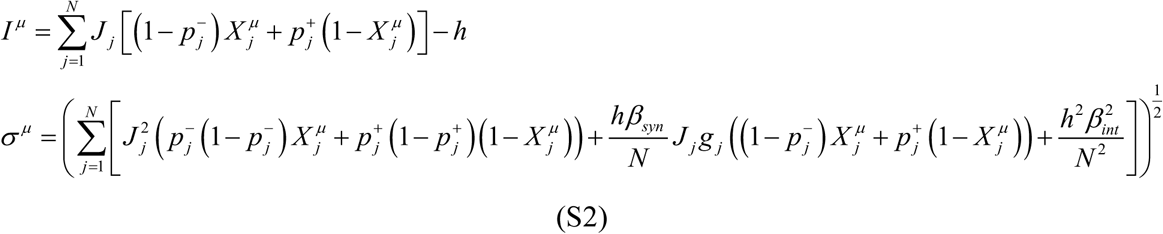

As a result, the inequality constraints on the probabilities of output spiking errors [line five in Eqs. (S1)] can be expressed in terms of *I*^*μ*^ and *σ*^*μ*^ :

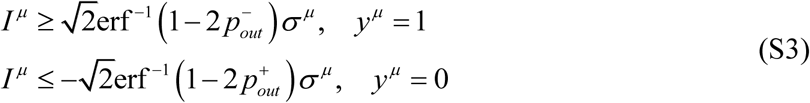

The above two inequalities can be combined into a single expression that must hold for a successfully learned association *μ*:

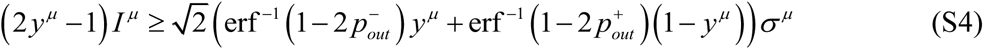

### C. Additional assumptions required for the replica calculation

Following the procedure outlined in (9, 26), we assumed that the model parameters 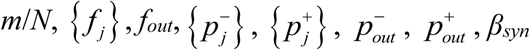 and *β*_*int*_ are intensive, or of order 1 with respect to *N*. In addition, we assumed that the connection weights are inversely proportional to the system size, 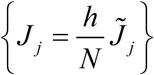, and refer to 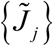 as scaled connection weights. This particular scaling is traditionally used in associative memory models (9), and it has been shown that in the biologically plausible high-weight regime, *Nwf ≫ h*, many model results become independent of this assumption (18). It follows from the second line of Eqs. (S1) that 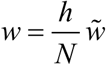, and we refer to 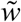 as scaled average that absolute connection weight.

The model, rewritten in terms of the scaled variables, contains one equality and *m* + *N* inequality constraints:

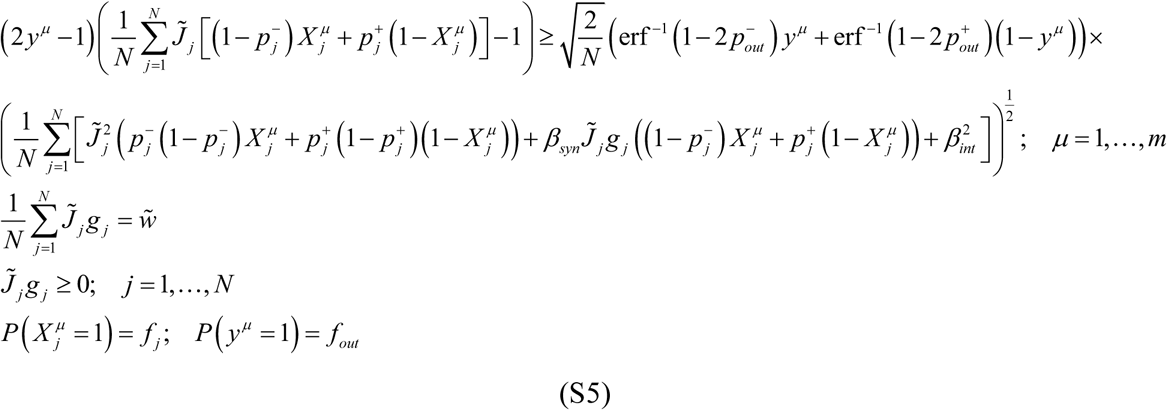

### D. Replica theory solution of the model

We begin by calculating the volume of the connection weight space, Ω({*X*^*μ*^, *y*^*μ*^}), in which Eqs. (S5) hold for a given set of associations, {*X*^*μ*^, *y*^*μ*^}:

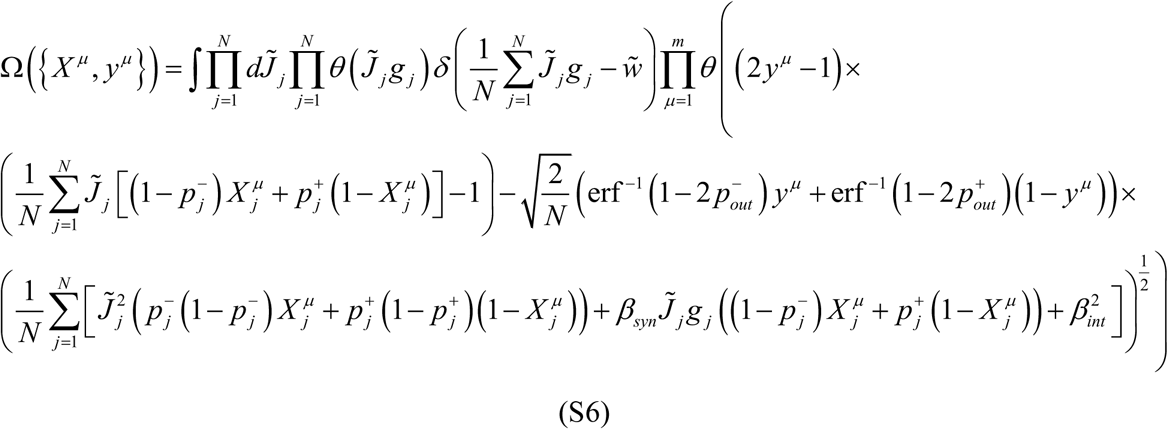

The typical volume of this solution space, Ω_*typical*_, is defined through the averaging of ln (Ω({*X*^*μ*^, *y*^*μ*^})) over the set of associations {*X*^*μ*^, *y*^*μ*^}, and is calculated by introducing *n* replica systems:

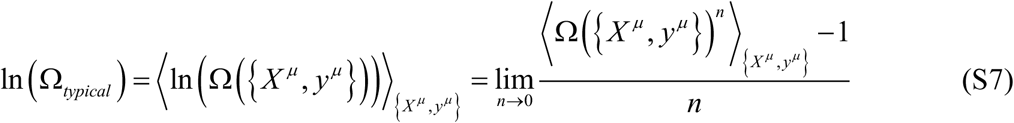

The quantity 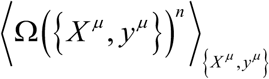 can be rewritten as a single multidimensional integral:

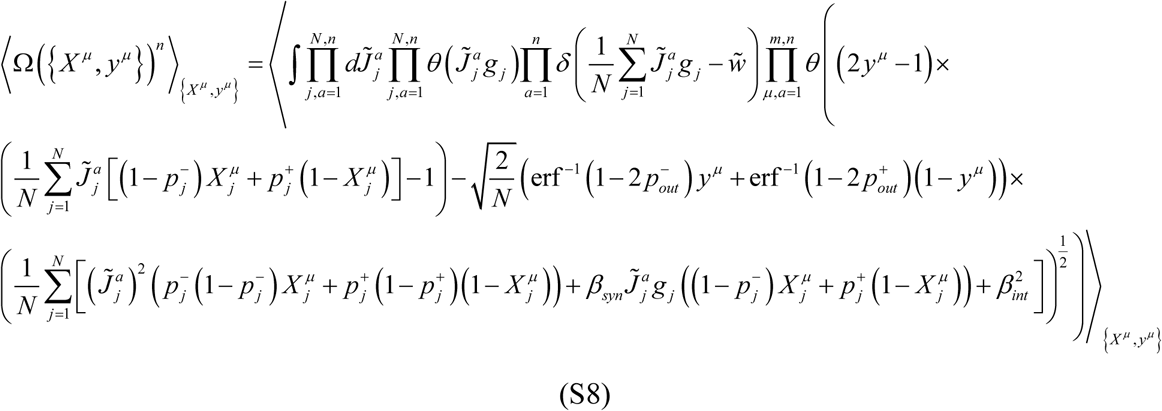

This integral was calculated by following a previously established procedure (9, 26), and below, we only outline the main steps of this calculation.

To calculate the average over the associations, we first decoupled *X*^*μ*^ and *y*^*μ*^ by introducing two new sets of variables:

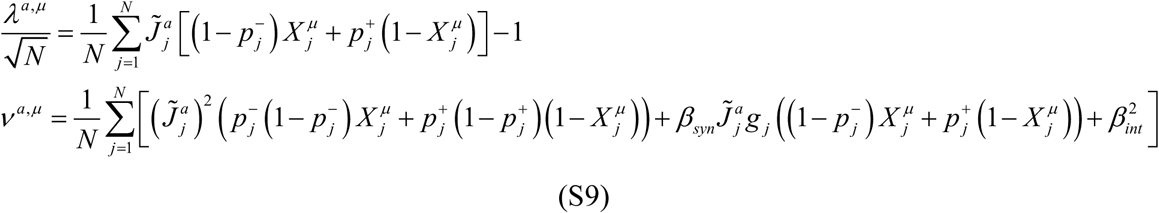

These variables were next incorporated into Eq. (S8) with the help of Dirac *δ*-functions:

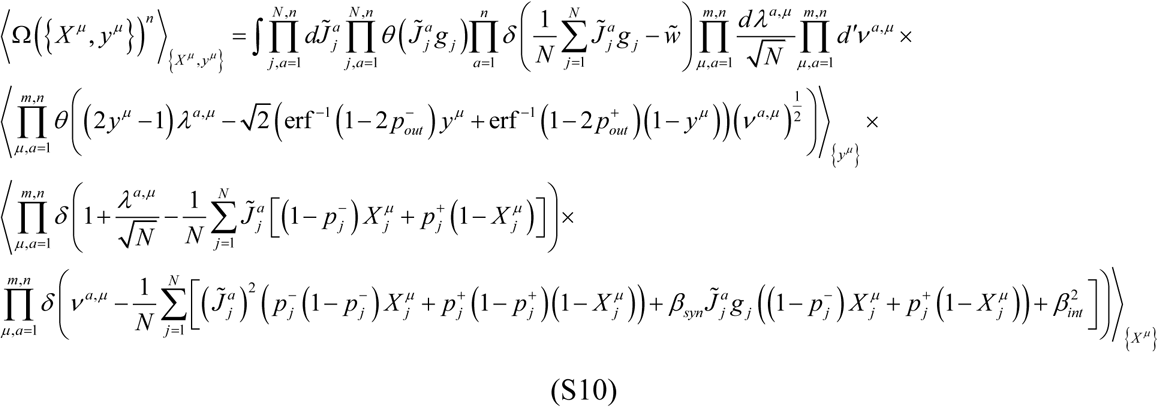

Symbol *d*′ in this expression and thereafter is designated for 0 to ∞ integration, whereas *d* is used for integration from -∞ to ∞. Next, the Heaviside step-functions and the *δ*-functions were replaced with their Fourier representations, making it possible to perform the averaging over the associations:

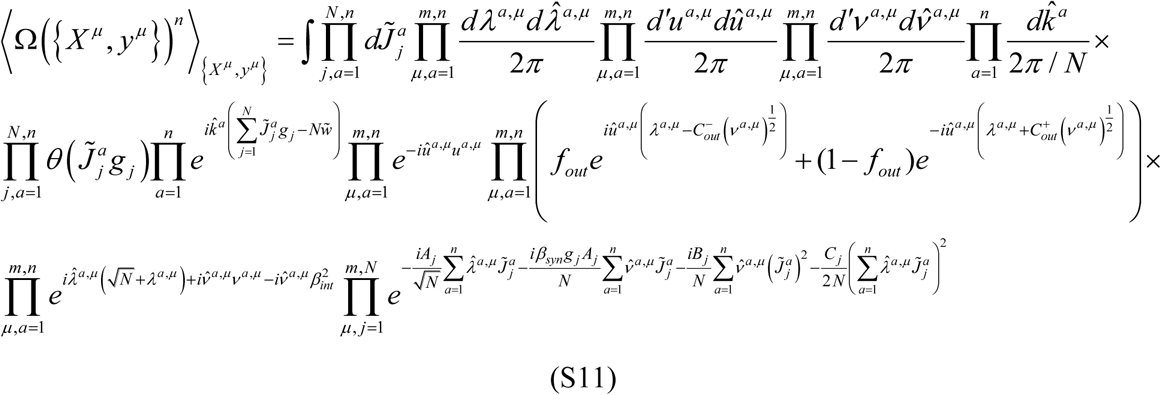

The following notation was used in the above expression:

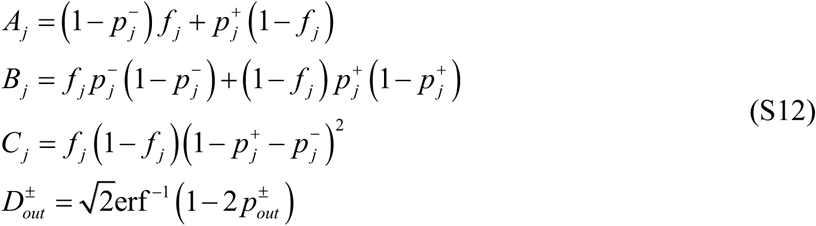

We note that parameters *A*_*j*_, *B*_*j*_, and *C*_*j*_ are nonnegative.

Next, we decoupled the products containing indices *j* and *µ* by introducing three sets of order parameters and embedding them into Eq. (S11) with the help of Dirac *δ*-functions as was done in Eqs. (S10, S11):

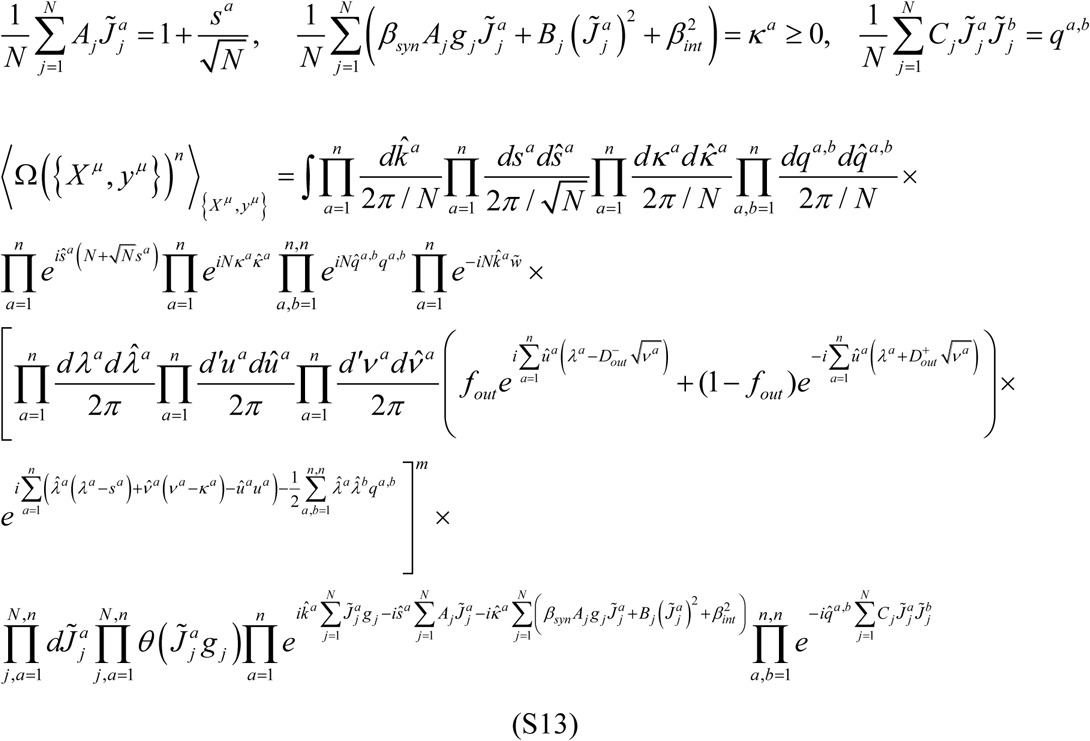

After integrating over 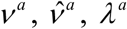 and 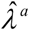 we obtained:

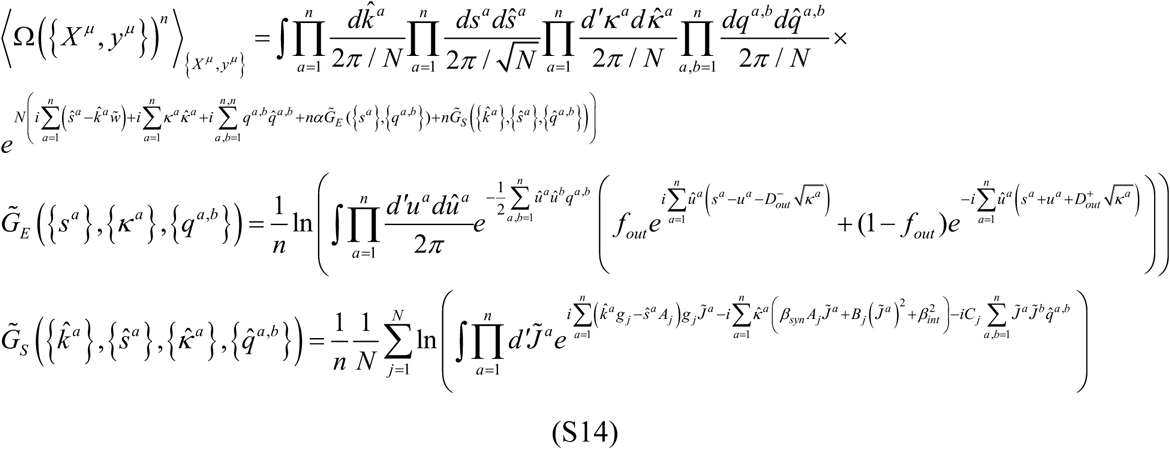

The above integral was calculated by using the steepest descent method (47) under the assumption of a replica symmetric saddle point, 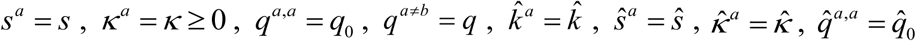 and 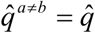

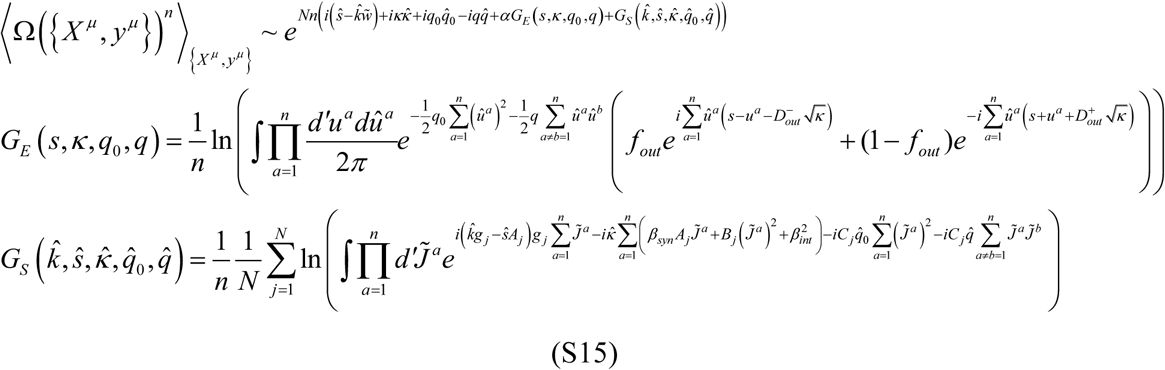

The non-redundant, replica symmetric saddle point coordinates 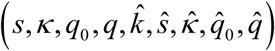 satisfy the following system of nine equations and one inequality:

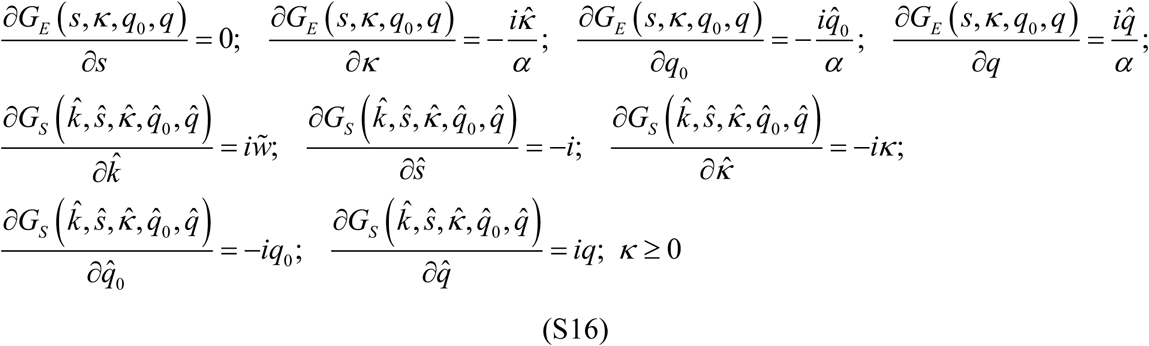

To simplify the expressions for *G*_*E*_ and *G*_*S*_ we employed the Hubbard-Stratonovich transformation [see e.g. (27)] and took the *n* → 0 limit:

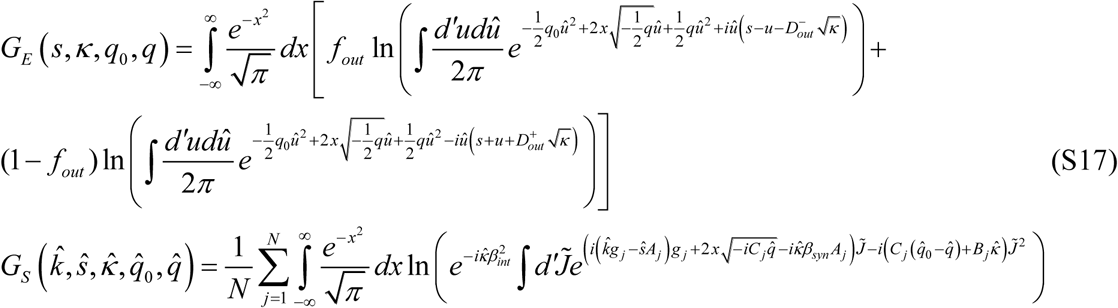

Integrals in the arguments of the natural logarithm functions were expressed in terms of the complementary error functions:

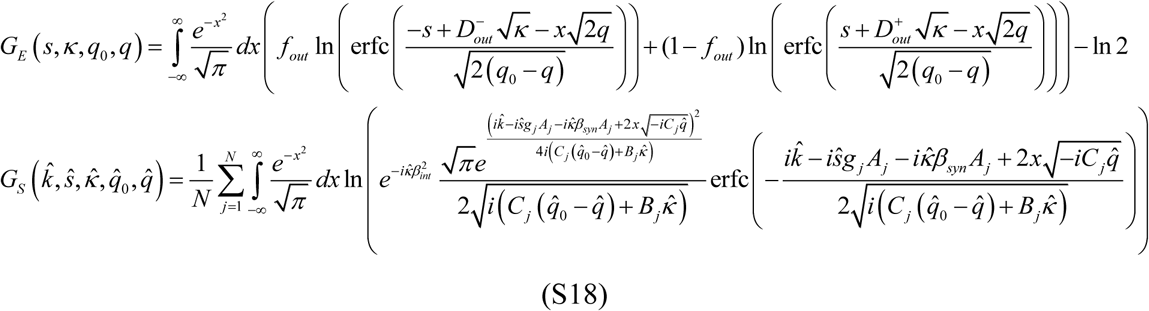

The following substitutions transform the replica symmetric saddle point coordinates into the real domain, 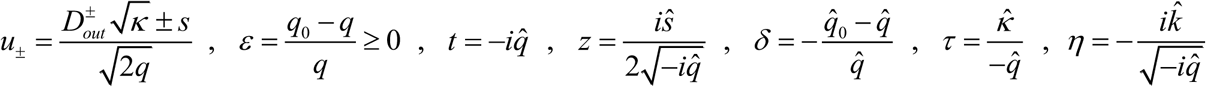, where these variables can be determined by solving the following problem:

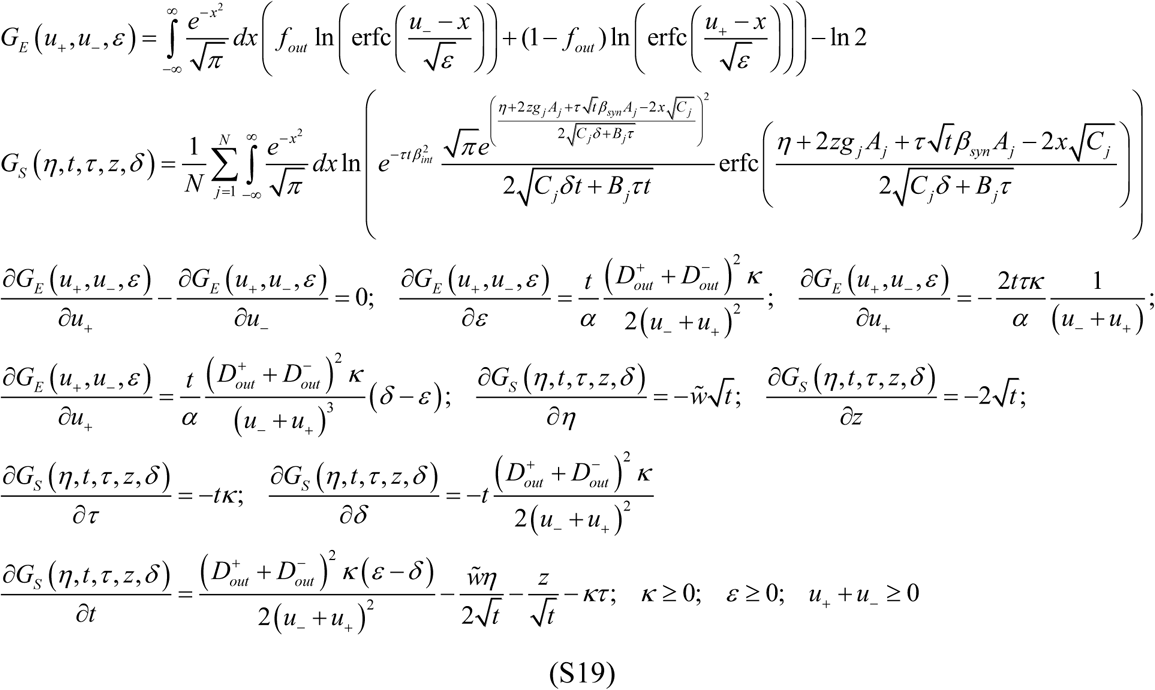

### E. Replica theory solution at critical capacity

With an increasing number of associations *m*, Ω_*typical*_ shrinks and approaches zero at the maximum (critical) capacity of the neuron, 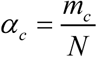. In this limit, (*q*_*0*_ − *q*) goes to zero and Eqs. (S19) can be expanded asymptotically in terms of 1/ *ε* and 1/ *δ*. In the leading order these expansions yield:

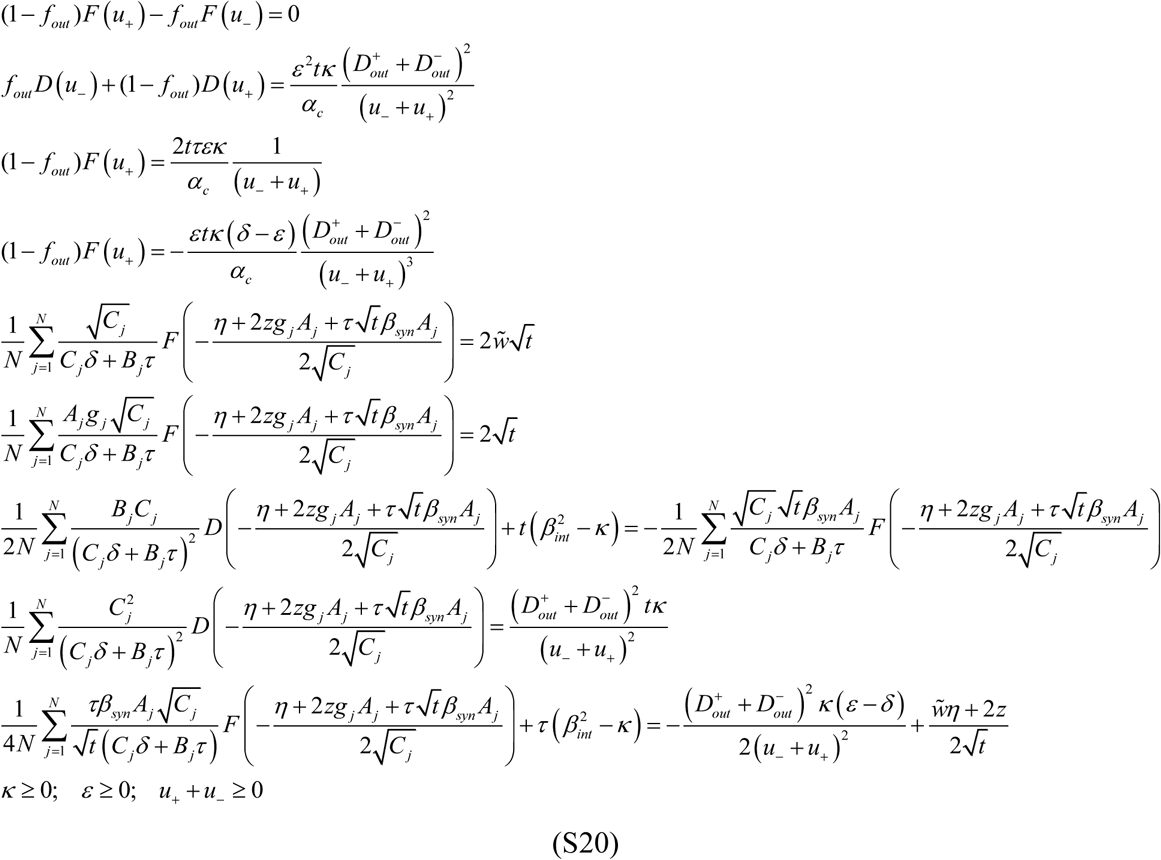

Functions *E, F*, and *D*, in Eqs. (S20) are defined as follows:

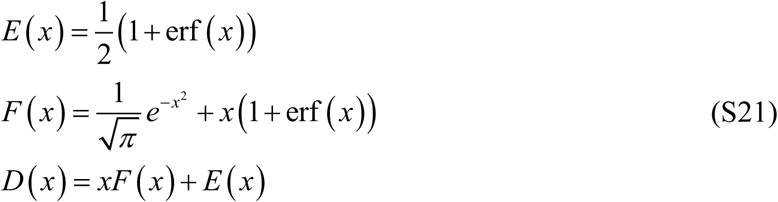

After replacing 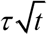 with *y, δ* / *τ* with *x*, and eliminating variables, *ε, t, κ, τ*, and *δ*, we arrived at the final system of six equations and one inequality. This system contains six latent variables *u*_±_, *x, η, y*, and *z* which determine the critical capacity of the neuron, *α*_*c*_ :

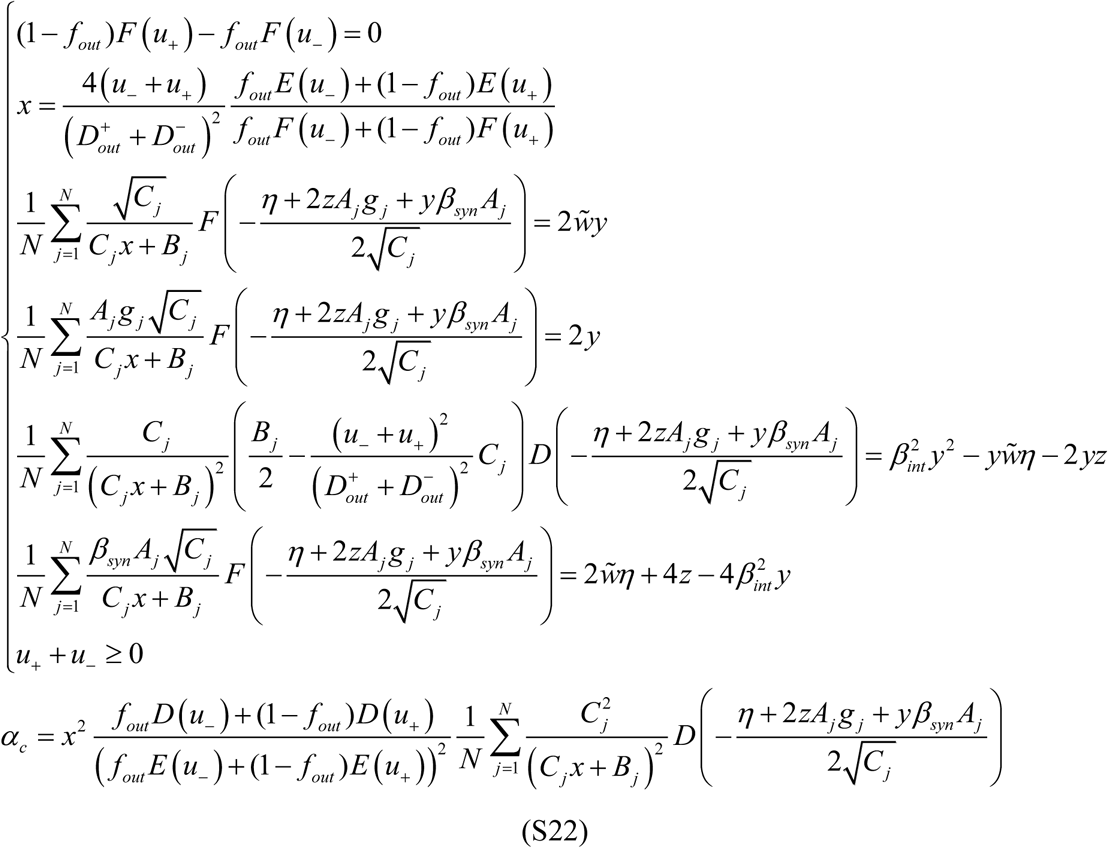

We note that Eqs. (S22) are in agreement with the solution described in (9), where a simplified version of the model presented here was solved by minimizing the probability of output spiking errors for a given intrinsic noise strength. Eqs. (S22) account for additional features such as the homeostatic constraint, learning by inhibitory inputs, heterogeneity of inputs, synaptic noise, and presynaptic spiking errors.

### F. Distribution of input weights at critical capacity

Connection probabilities, *P*^*con*^, and probability densities for non-zero input weights, *p*^*PSP*^, at critical capacity were calculated as previously described (9, 26). The result depends on the set of latent variables of Eqs. (S22):

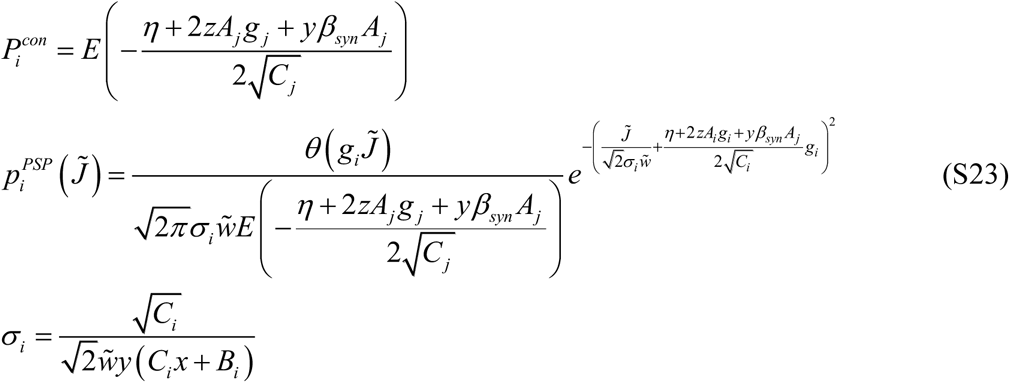

A given input, *i*, has a non-infinitesimal probability of having a connection weight of zero, while its probability density for having non-zero connection weights is a truncated Gaussian with a standard deviation 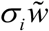.

### G. The solution in the case of balanced spiking errors

Without loss of generality, we assumed that the expected rates of erroneous spikes and spike failures are equal, and, as a result, these spiking errors do not affect the input and output firing probabilities {*f*_*j*_} and *f*_*out*_ :

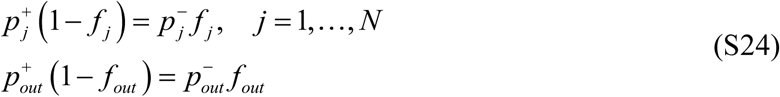

In this case, it is more convenient to express the results of the model in terms of spiking error probabilities:

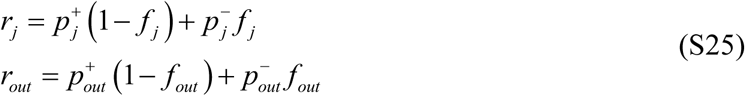

This change does not alter Eqs. (S22, S23), but the coefficients defined in Eqs. (S12) transform into:

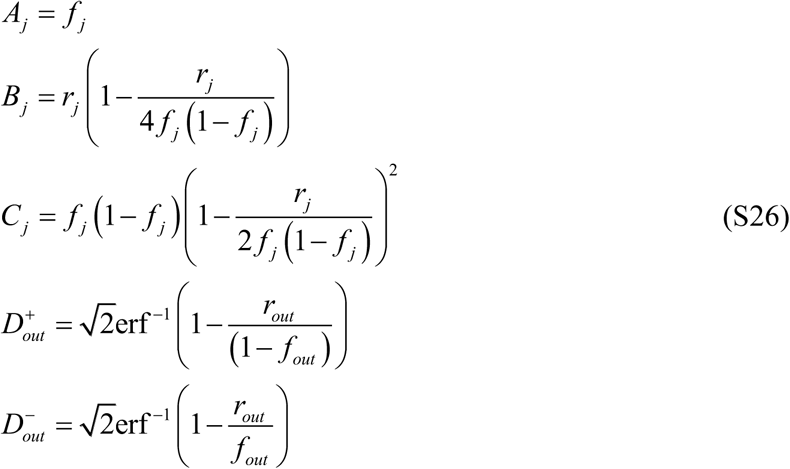

Eqs. (S22, S23) were solved in MATLAB to produce the results for heterogeneous networks consisting of inhibitory and excitatory neurons with distributed spiking error probabilities (Figures 7A-C of the main text) and distributed intrinsic and synaptic noise strengths (Figures 7D-F of the main text). In both cases, the remaining model parameters were the same for all neurons. We note that in both cases the solutions of Eqs. (S22, S23) depend on *β*_*int*_ and *β*_*syn*_ only in a combination 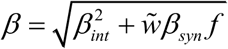. This parameter is referred to as the postsynaptic noise strength.

The code is available at https://github.com/neurogeometry/Associative_Learning_with_Noise.

### H. The solution in the case of two homogeneous classes of inputs and balanced spiking errors

In this case, all neuron inputs have the same firing probability, *f*, and the same spiking error probability, *r*. Eqs. (S22, S23) simplify significantly after the introduction of two new variables, 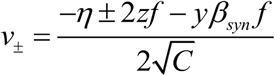:

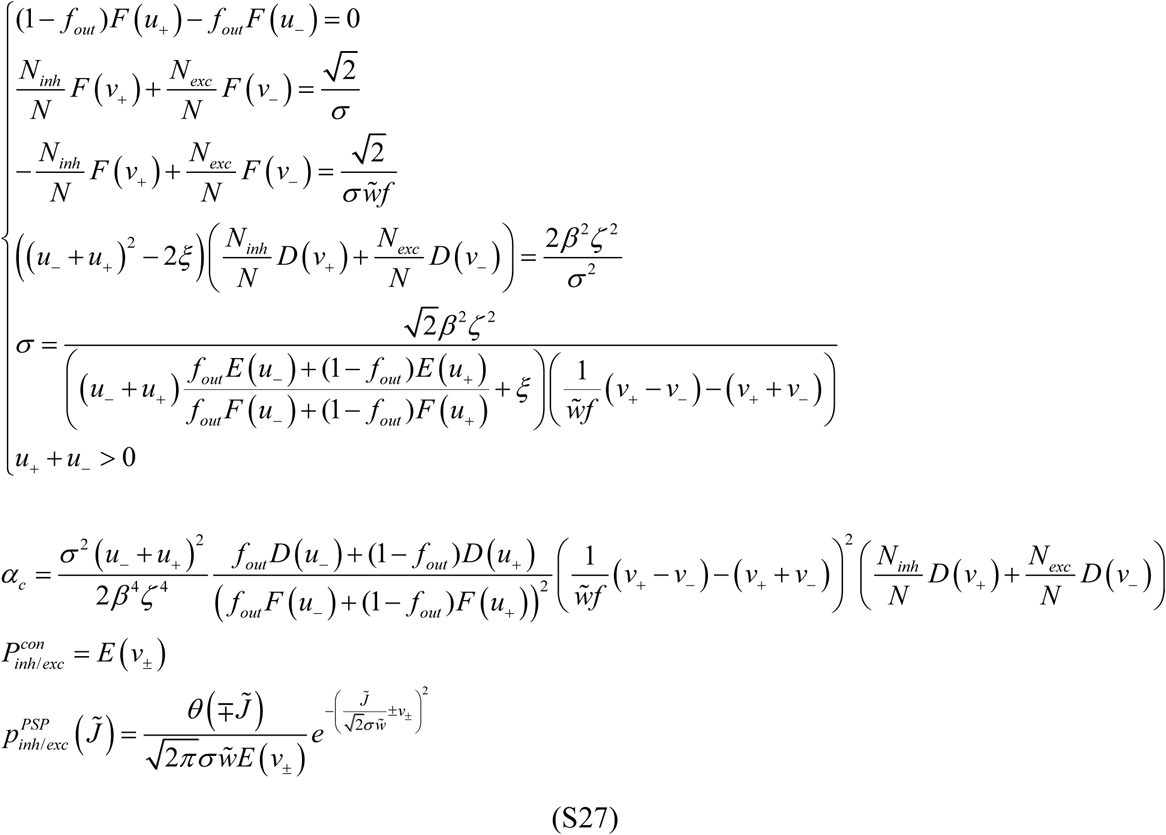

The intrinsic and synaptic noises in Eqs. (S27) are entirely contained within the parameter *β*, while the spiking error probabilities *r* and *r*_*out*_ appear only in the parameters *ξ* and *ζ*:

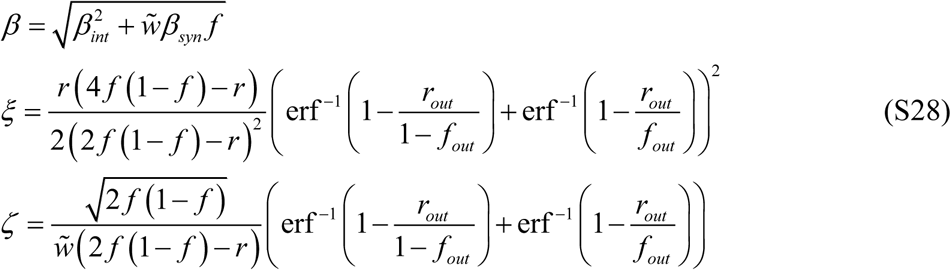

We note that in the absence of spiking errors in the input (*r* = 0), Eqs. (S27) are similar in structure to the solution of a simplified model considered by us previously (18). That model did not explicitly consider different sources of errors and noise, but instead, used a generic parameter *ρ*, referred to as the rescaled robustness, to ensure that memories are recalled reliably in the presence of intrinsic noise only. Solutions to both models become identical when *ρ* = *βζ*.

For associative networks considered in the main text, we set *r*_*out*_ = *r* and *f*_*out*_ = *f* in Eqs. (S27, S28) as these parameters in the homogeneous case must be the same for all neurons in the network. Figures 1, 4, and 6 of the main text show the results of the homogeneous model as functions of *β* and *r*.

### I. Numerical solution of the model with nonlinear optimization

For a finite number of inputs, the solution to the problem outlined in Eqs. (S5) was obtained numerically. To that end, we made the problem feasible by introducing a slack variable *s*^*μ*^ ≥ 0 for every association and chose the solution that minimizes the sum of these variables:

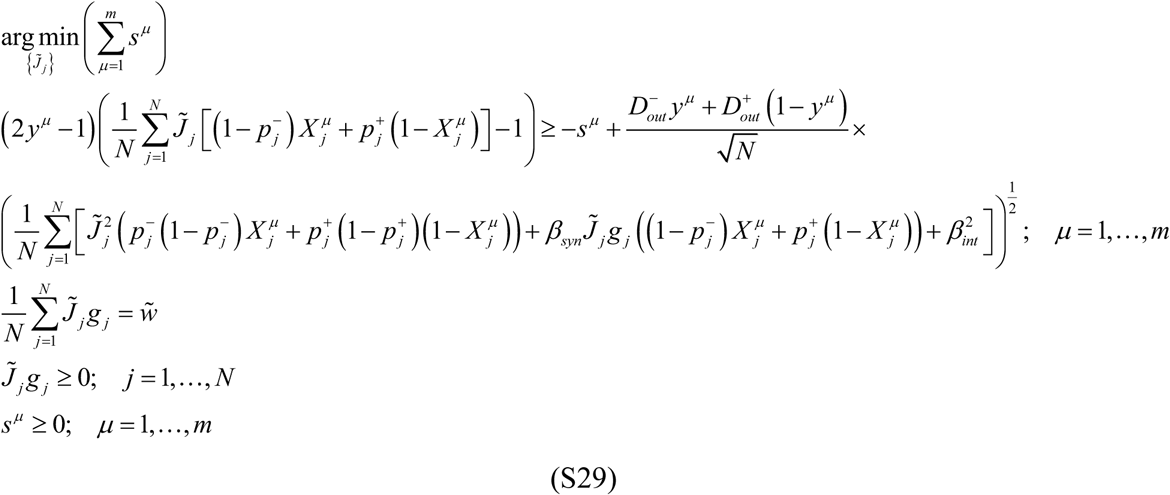

Eqs. (S29) were solved by using the *fmincon* function of MATLAB and the results are shown in Figures 2, 3, 5, and 6 of the main text. The *fmincon* function utilizes the interior-point technique for finding solutions of constrained nonlinear optimization problems (48, 49). The code is available at https://github.com/neurogeometry/Associative_Learning_with_Noise.

### J. Numerical solution of the model with a perceptron-type learning rule

In addition to the replica and nonlinear optimization solutions, a biologically more plausible online solution of Eqs. (S29) was devised by approximately stepping in the direction of the negative gradient of the sum of the slack variables. The latter is,

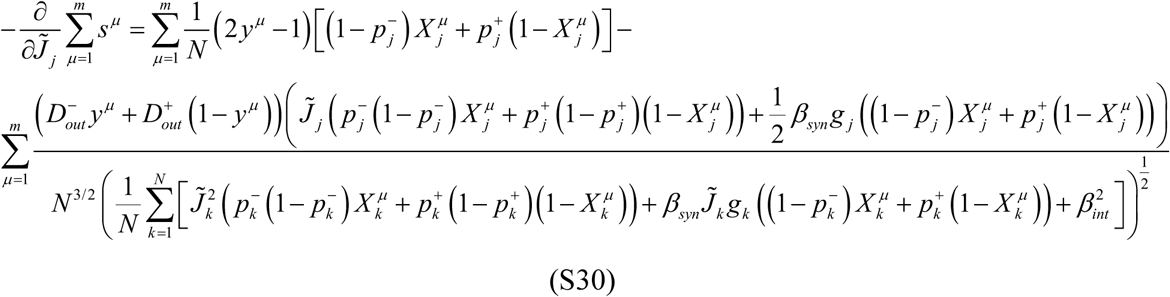

The first approximation to this gradient was made by ignoring the second term in the r.h.s. of Eq. (S30). This was done because there is no clear way of calculating this term in an online, biologically plausible manner. The second approximation was made by noting that 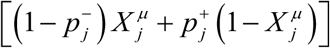 in the r.h.s. of Eq. (S30) is the average of 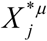 over the spiking errors, and therefore, a stochastic estimate of this gradient direction can be made in an online manner with a perceptron-type learning step 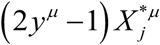 (45).

With these approximations, we proposed the following learning rule. At each learning step, e.g. *μ*, a neuron receives an input containing spiking errors, *X*^\**μ*^, combines it with its synaptic and intrinsic noise, and produces an output, *y*^\**μ*^. If this output differs from the neuron’s target output, *y*^*μ*^, its input connection weights are updated in four consecutive steps:

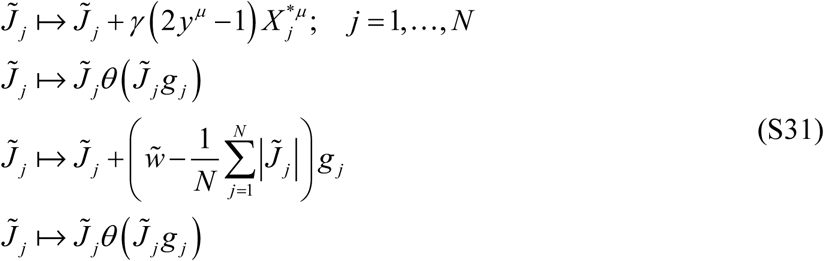

By including presynaptic spiking errors, synaptic and intrinsic noise in the condition that triggers the learning steps outlined in Eqs. (S31), the learning rule implicitly depends on the model parameters {*r*_*j*_}, {*β*_*syn, j*_}, and *β*_*int*_ describing the neuron’s intrinsic noise. By approximately minimizing the sum of the slack variables, the learning rule minimizes the neuron’s output spiking error probability for a given memory load, and when the feasible solution of Eqs. (S29) is found (i.e. *s*^*μ*^ = 0; *μ* =1,, *m*), the neuron’s output spiking error probability falls below the desired probability, *r*_*out*_ (Figure 6A). At capacity, the two probabilities must equal and, therefore, the learning rule also depends implicitly on *r*_*out*_.

Unlike the standard perceptron learning rule, Eqs. (S31) are based on noisy input and enforce sign and homeostatic constraints during learning. The first update step in Eqs. (S31) is a stochastic perceptron learning step, in which parameter *γ* is referred to as the learning rate. The second step is introduced to enforce the sign constraints, while the last two steps combined implement the homeostatic *l*_1_-norm constraint and are equivalent to the soft thresholding done in LASSO regression (46). A related rule, in the absence of errors, noise, and *l*_1_-norm constraint, was previously described in (9, 50). In numerical simulations, we trained neurons on associations presented in the order of their appearance in the associative sequence, one at a time. This constitutes one learning epoch. We set *γ* = 0.1 and ran the algorithm until a solution was found or the maximum number of 10^6^ epochs was reached. The results of this procedure are shown in Figure 6 of the main text.

### K. Mutual information contained in retrieved associative sequences

The mutual information contained in one successfully retrieved association (*X*^*μ*^ → *X*^*μ* +1^) can be calculated as a difference of marginal and conditional entropies,

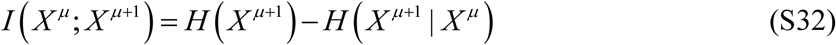

For homogeneous networks loaded with associations consisting of random and independent network states with balanced spiking errors the two entropies reduce to:

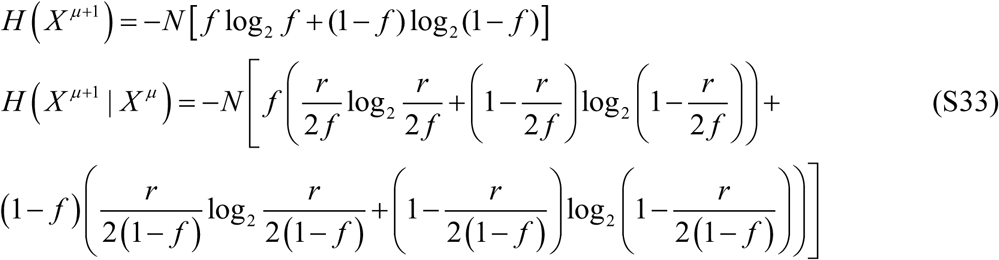

As the length of a retrieved sequence may be shorter than the length of the loaded sequence, *m*, in the main text we considered two types of retrieved information. One type is defined as the expected retrieved information per memory playout in which contributions of partially retrieved sequences are set to zero. This information is based on completely retrieved sequences only and is equal to the product of the retrieval probability (Figure 2C) and *mI*. The other type of retrieved information is calculated based on completely and partially retrieved sequences and is equal to the product of the average retrieved sequence length and *I*. According to these definitions, the former is always less or equal to the latter.

